# Time-Restricted Feeding Extends Healthspan in Both Sexes and Lifespan in Male C57BL/6J Mice

**DOI:** 10.1101/2025.10.22.683527

**Authors:** Samantha E. Iiams, Nathan J. Skinner, Mary Wight-Carter, Victoria A. Acosta-Rodríguez, Carla B. Green, Joseph S. Takahashi

## Abstract

Time-restricted feeding (TRF) aligned with an organism’s circadian rhythm has been shown to improve health, but its long-term effects on healthspan and lifespan in mammals, especially under normal dietary conditions, remain unclear. Here, we examined the impact of 12-hour (h) and 8h nightly TRF windows in male and female mice fed regular chow. TRF improved multiple health measures, including behavioral rhythmicity, body weight and composition, frailty, and disease onset. These effects were most pronounced in the 8h-TRF group, which exhibited voluntary caloric restriction in addition to time restriction. A composite Healthspan Index revealed that TRF extended healthspan in both sexes, though the benefits were more prolonged in females relative to their total lifespan. Median lifespan was significantly extended in males under 8h-TRF by 12%, whereas females showed no significant lifespan extension, highlighting sex-specific responses to TRF.

## Main Text

While human life expectancy has nearly doubled since the early 20th century (*1*), healthspan remains limited, with many spending over 10% of their lives in poor health due to age-related functional declines and disease development (*2*). Caloric restriction (CR), a potent aging intervention involving a 30-50% reduction in food intake without malnutrition, has been extensively studied for its ability to delay the onset of age-associated diseases, including cancer, cardiovascular disorders, and neurodegeneration, while extending lifespan across multiple model organisms (*3–5*). Incorporating daily feeding/fasting cycles with CR enhances these geroprotective effects (*6–9*), and aligning feeding with an organism’s active circadian phase can further extend lifespan (*10*). This synergy likely reflects both the synchronization of nutrient intake with internal metabolic processes and the protective effects of temporal compartmentalization, driven by circadian clock-regulated gene expression in the liver and other peripheral tissues (*11–18*).

Aging triggers a progressive damping of circadian rhythmicity in both humans and animal models. This decline, evident in behavior and physiology, stems from a reduction in amplitude and shifted phases of the molecular clock’s transcriptional output (*19*). Interventions that restore circadian rhythms can improve health and extend longevity in model organisms (*20*), suggesting that optimizing meal timing is a promising strategy to counteract age-related rhythm disruptions and promote healthy aging (*21*).

Given that CR is challenging to maintain in people (*22*), meal timing alone via time-restricted feeding (TRF) in animals, or time-restricted eating (TRE) in humans, has emerged as an alternative strategy to delay aging (*23, 24*). TRF, which restricts feeding to specific daily windows without limiting overall caloric intake, can confer similar benefits as timed CR, including improved rhythmic gene expression, cognitive function, and cardiometabolic health, even in obesity models fed a high-fat, high-sucrose diet (*25–32*). Recent clinical TRE trials report high adherence and improved cardiometabolic outcomes in subjects with pre-existing conditions, demonstrating its potential as a feasible intervention to enhance human healthspan (*33–37*).

However, few studies have examined the benefits of TRF/TRE under typical metabolic conditions without diet– or genetically-induced obesity (*38–40*), and even fewer have compared impacts between the sexes (*41*). The long-term impact of TRF on healthspan and lifespan in mammalian models also remains unclear (*42, 43*). In this study, we sought to address these key knowledge gaps and determine whether TRF could serve as an effective aging intervention for healthy male and female mice.

## Results

### Early Onset Circadian Aligned TRF Has Sex-Specific Impacts on Longitudinal Feeding and Wheel-Running Activity Levels

To determine how time-restricted feeding (TRF) impacts healthspan and lifespan in mammals, we individually housed 264 female and 264 male C57BL/6J mice in wheel cages with automated feeders at 2 months of age. To measure impacts under normal nutritional conditions, mice were fed purified precision pellets (Bioserv F0075). For the first 8-weeks, all mice were fed *ad libitum* (free-feeding; AL) with access to a maximum of 22 pellets or 6.6g (23.8 Kcal) per day. At 4 months of age, N=78/sex were switched to a 12-hour (h) TRF window (Zeitgeber time (ZT) 12-24) and another N=78/sex to an 8h-TRF window (ZT14-22) both circadian aligned to the night, the active phase of nocturnal rodents. The remaining N=108/sex were maintained AL to serve as the control. Across all feeding groups, mice were provided with a daily allotment of food that exceeded their typical intake, and no group ever consumed the full amount, ensuring that no caloric restriction was imposed (Fig. 1, A and B, and Fig. S1) (*44*). These feeding regimens were maintained for the duration of the study. Any food hoarded within the cage was removed and excluded from Fig. 1B and Figs. S2-3 as described in the methods.

**Figure 1:**
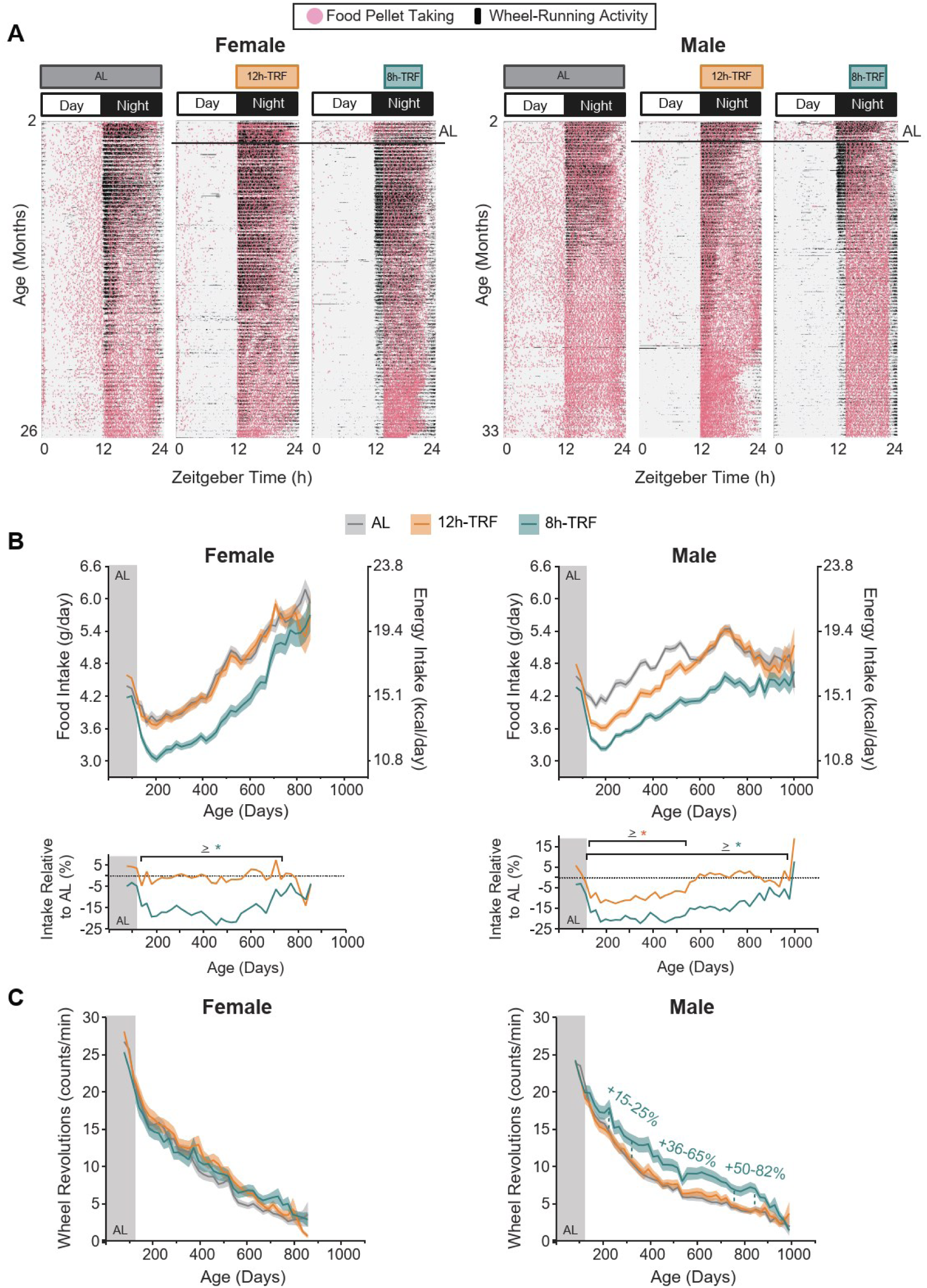
Circadian aligned TRF reduces feeding and improves wheel-running activity in a feeding window– and sex-dependent manner. (**A**) In females 2-26 months of age and males 2-33 months of age, representative actograms of individual mice show wheel-running (black mark) and food pellet taking (pink dot) activity over the course of the 24-hour (h) light:dark cycle. Black horizontal line shows ends of *ad libitum* (AL) baseline feeding for the TRF groups at 4 months of age. (**B**) (*Top*) Longitudinal profiles of average daily food intake and energy intake with age and (*Bottom*) percent differences in CR relative to AL controls with age. Two-way ANOVA: Females: Age=****, Feeding=****, Interaction=****; Males: Age=****, Feeding=****, Interaction=**** (Full details of ANOVA results for each figure can be found in Table S1). (**C**) Longitudinal profiles of average daily wheel revolutions with age. Two-way ANOVA: Females: Age=****, Feeding=**, Interaction=ns; Males: Age=****, Feeding=****, Interaction=ns. <u>B and C:</u> AL baseline feeding (Gray area). TRF feeding conditions (White area). Each point represents mean of 21 days ± SEM (shaded regions). Asterisks/percents represent significant differences in TRF vs AL (*P ≤ 0.05; **P ≤ 0.01; ***P ≤ 0.001; ****P ≤0.0001) as determined by two-way ANOVA and Tukey’s post-hoc. Females N=71-103 per group. Males N=71-97 per group.

### Food Intake

Measurements of daily food intake throughout each mouse’s lifespan showed that feeding increased over time in both sexes, then declined near the end of life in males, similar to previous mouse studies (Fig. 1, A and B, and Fig. S2-3) (*10, 44*). In line with other rodent longitudinal recordings (*45*), females in our study consumed more food relative to their body weight at older ages than males. We also observed sex-specific responses to TRF where 12h-TRF females consumed the same amount as AL, while 12h-TRF males significantly reduced intake compared to AL by 8-14% from 140-539 days (∼5-18 months) (Full details of ANOVA results for all figures can be found in Table S1). In contrast, 8h-TRF self-imposed caloric restriction (CR) for the majority of life in both sexes, with intake in females reduced by 10-22% between 140-728 days (∼5-24 months) and males reduced by 9-23% between 140-980 days (∼5-33 months) relative to sex– and age-matched AL controls. This 8h-TRF schedule, which narrows the daily eating window without explicitly limiting calorie intake as seen in prior mouse and human studies (*46, 47*), presents an alternative strategy for achieving long-term CR in both sexes.

Although actograms (Fig. 1A) show occasional pellet-taking events outside the defined TRF windows, these largely reflect a technical limitation of the feeding system: pellets dispensed at the very end of the dark phase cannot be retracted, allowing a single pellet to remain available and be retrieved during the light period. Importantly, this represents at most one 300 mg pellet per day, accounting for ≤10% of daily intake in 8h-TRF females (lifetime intake 3.0–5.7 g), ≤8% in 12h-TRF females (3.6–5.9 g), ≤9% in 8h-TRF males (3.2–4.7 g), and ≤8% in 12h-TRF males (3.6–5.4 g). We quantitatively assessed this behavior across the full lifespan and found the true average proportion of pellets taken outside the feeding window was low, ranging from 5.17–6.34% across feeding groups and sexes (Fig. S1B). In addition, food hoarded within the cage accounted for no more than ∼10–12% of the amount removed from the feeder (Fig. S1C). Together, these data suggest that the majority of food intake in both TRF schedules occurs within the intended feeding window. In contrast, some daytime pellet-taking in AL mice is expected, as C57BL/6J mice naturally consume a portion of their food during the inactive phase, with ∼75% of intake occurring at night under AL conditions (*9, 10*).

### Wheel-Running

Analysis of daily wheel-running recordings for each mouse revealed that individuals in all feeding groups, of both sexes, consistently displayed nocturnal activity, which subsequently declined with age, as anticipated (Fig. 1, A and C, and Figs. S4-5) (*48, 49*). Only 8h-TRF males had increased levels of wheel activity from 224-833 days of age (∼7-28 months) with the enhanced activity ranging +36-82% from 329-833 days (∼11-28 months) relative to AL. This male-specific difference further underscores the sex-specific effects of TRF on behavior. Additionally, in line with previous work (*10, 50*), the sustained increases in activity under 8h-TRF suggest a healthspan-enhancing effect in male mice.

### TRF Improves the Amplitude of Daily Rhythmic Behaviors

We next examined age-related changes in the diurnal profiles of feeding and wheel-running, two well-established rhythmic behaviors, as high-amplitude behavioral rhythms are closely associated with improved health outcomes (Fig. 2, A and B) (*19, 20, 51, 52*). As intended, TRF reduced the feeding window, shown by longitudinal measurements of the time required for each group to retrieve 90% of their daily food intake from the feeder. Across both sexes, mice under 12h-TRF took 90% of their food within 12h, whereas those under 8h-TRF did so within 8h (Fig. 2C). The time required to reach this threshold declined with age across groups, particularly in females, indicating that feeding became more consolidated in older animals (Fig. 2, A and C). Consistent with this pattern, both TRF groups showed significantly longer daily maximum fasting durations than AL controls, exceeding 10h in the 12h-TRF group and 14h in the 8h-TRF group across both sexes (Fig. 2D).

**Figure 2:**
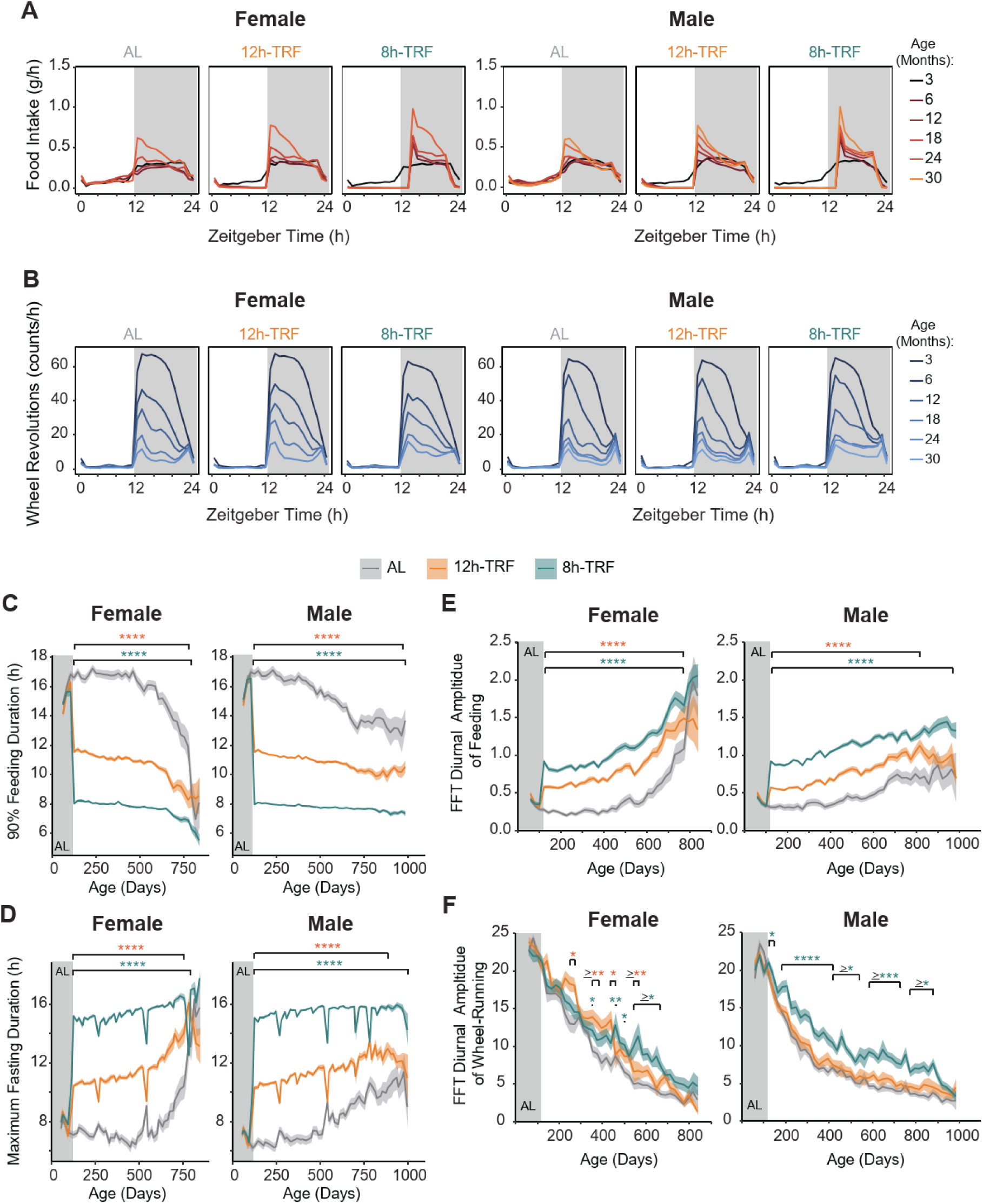
TRF improves the amplitude of daily feeding and wheel-running rhythms. (**A**) 24h profiles of feeding and (**B**) wheel-running starting at 3 and then every 6 months of age. 12h light (white area), 12h dark (gray area). (**C**) Longitudinal plots for daily time it takes each group of mice to take 90% of their daily intake from the feeder. Type III ANOVA (Wald χ²): Females: Age=****, Feeding=****, Interaction=****; Males: Age=****, Feeding=****, Interaction=****. (**D**) Longitudinal plots for daily maximum fasting times achieved. Type III ANOVA (Wald χ²): Females: Age=****, Feeding=****, Interaction=****; Males: Age=****, Feeding=****, Interaction=****. To note: Large, transient dips in fasting due to interruptions in data acquisition (e.g., fasting blood collection attempt and computer outage). (**E**) Longitudinal FFT plots of the diurnal amplitude of feeding with age. Type III ANOVA (Wald χ²): Females: Age=****, Feeding=****, Interaction=****; Males: Age=****, Feeding=****, Interaction=****. (**F**) Longitudinal FFT plots of the diurnal amplitude of wheel-running with age. Type III ANOVA (Wald χ²): Females: Age=****, Feeding=*, Interaction=****; Males: Age=****, Feeding=****, Interaction=***. <u>C-F:</u> AL baseline feeding (Gray area). TRF feeding conditions (White area). Each point represents mean of 14 days ± SEM (shaded regions). Asterisks represent significant differences in TRF vs AL (*P ≤ 0.05; **P ≤ 0.01; ***P ≤ 0.001; ****P ≤0.0001) as determined by type III ANOVA (Wald χ²) and Holm’s post-hoc. <u>For all panels</u>: Females N=71-103 per group. Males N=71-97 per group.

A Fast Fourier Transform (FFT) analysis of diurnal amplitude, reflecting the strength of daily rhythms, confirmed that TRF increased feeding amplitude and led to a preservation of enhanced rhythmicity with age, particularly in the 8h-TRF group. (Fig. 2, A and E). While the diurnal amplitude of wheel-running declined with age across all groups, 8h-TRF males maintained stronger rhythms throughout life compared to controls (Fig. 2, B and F). In females, TRF modestly increased wheel-running amplitude, with intermittent enhancements relative to controls from ∼249 days onward and a sustained elevation in the 8h-TRF group between 543–669 days of age. As mice age, TRF, particularly an 8h window, counteracted the natural decline in behavioral rhythmicity by consolidating food intake and preserving higher-amplitude wheel running activity compared with AL controls.

### TRF Improves Body Composition with Limited Effects on Systemic Metabolic Markers

#### TRF improves body weight and composition

Body weights were recorded every 21 days (Fig. 3A, and Fig. S6), and body composition was analyzed every 6 months using nuclear magnetic resonance imaging (EchoMRI, Houston, TX) (Fig. 3, B and C). In females, 12h-TRF without CR lessened age-related gains in body weight by 5-8% from 350-686 days of age (∼12-23 months) compared with AL controls. This feeding restriction also improved fat vs lean body composition, reducing fat mass while increasing lean mass by 3-4% at 12 and 18 months of age. Unexpectedly, 8h-TRF females with self-imposed CR did not exhibit any further improvements beyond 12h-TRF, in weight or fat/lean composition. In males, 12h-TRF reduced body weight gain by 5-7% from 224-476 days (∼7-16 months) and improved fat vs lean composition by 3% at 12 months. 8h-TRF in males enhanced these benefits, lowering body weight gain by 5-16% from 161-749 days (∼5-25 months) and improving fat vs lean composition by 3-7% from 6-12 months. Ultimately, these results demonstrate that while 12h-TRF is sufficient to attenuate weight gain and optimize body composition in both sexes, the benefits are more prolonged in females; conversely, males derive significantly greater long-term physiological advantages from a more restrictive 8h-TRF schedule combined with self-imposed CR.

**Figure 3:**
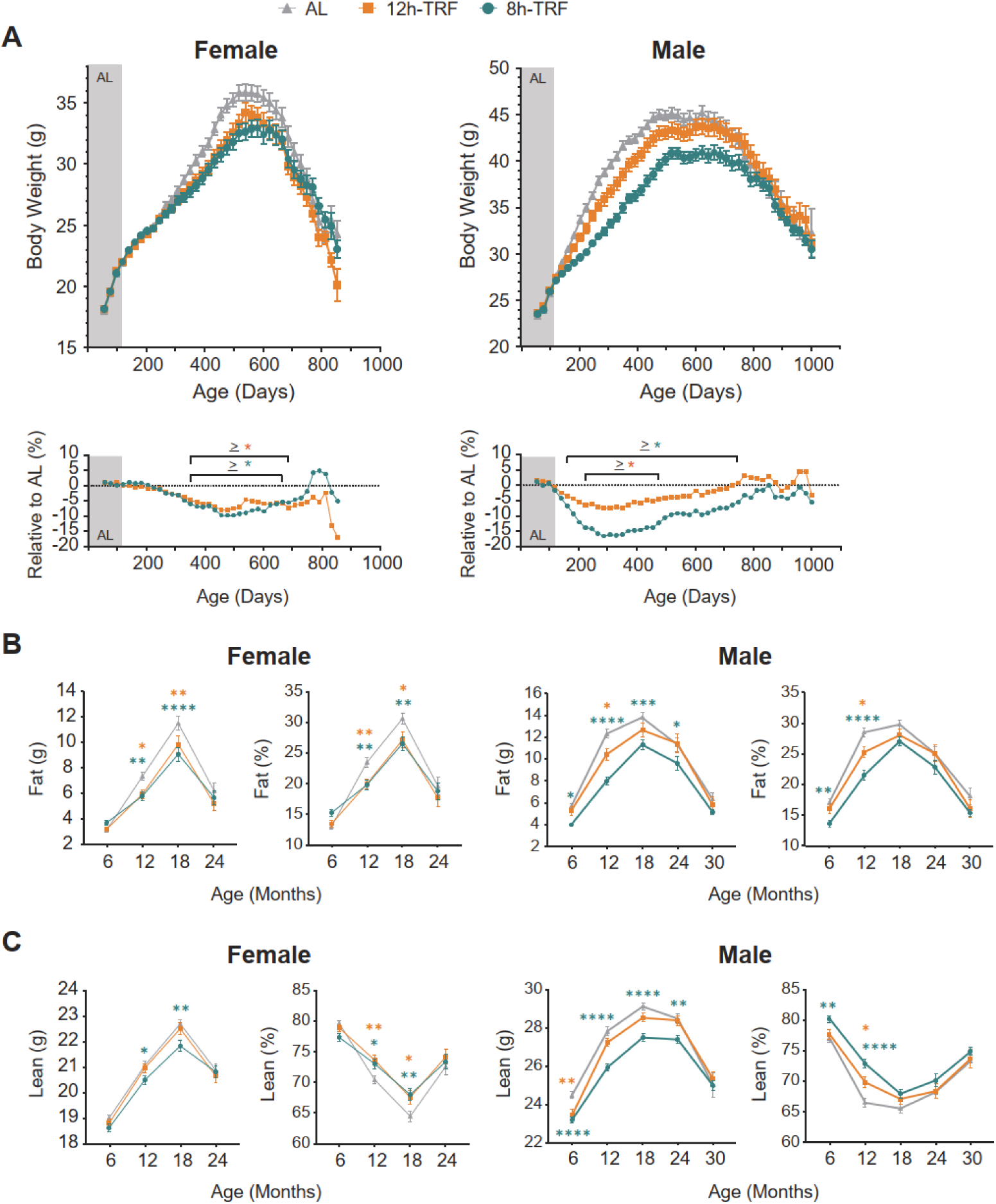
A 12h TRF window is sufficient to reduce body weight and fat mass in both sexes. (**A**) (*Top*) Longitudinal profiles of body weight and (*Bottom*) percent differences in body weight relative to AL controls with age. Two-way ANOVA: Females: Age=****, Feeding=****, Interaction=****; Males: Age=****, Feeding=****, Interaction=****. AL baseline feeding (Gray area). TRF feeding conditions (White area). Each point represents mean of 21 days (cage change) ± SEM (bars). Asterisks represent significant differences in TRF vs AL (*P ≤ 0.05; **P ≤ 0.01; ***P ≤ 0.001; ****P ≤0.0001) as determined by two-way ANOVA and Tukey’s post-hoc. Females N=76-103 per group. Males N=76-102 per group. (**B**) Total fat mass as a percentage of body weight with age. (**C**) Total lean mass as a percentage of body weight with age. <u>B and C:</u> Means every 6 months ± SEM (bars). Asterisks represent significant differences in TRF vs AL as determined by two-way ANOVA and Tukey’s post-hoc. Females 6-24 months: AL, N=43-101. 12h-TRF, N=32-71. 8h-TRF, N=38-72. Males 6-30 months: AL, N=23-102. 12h-TRF, N=19-77. 8h-TRF, N=36-75.

#### TRF reshapes diurnal RER and EE rhythms

Metabolic changes under TRF were assessed using indirect calorimetry at 19 months of age. Mice were randomly selected and individually housed in Promethion metabolic cages equipped with automated feeders for 5-6 days at room temperature (25°C), followed by 5 days at thermoneutrality (30°C), to assess metabolic parameters independent of the energetic demands of thermoregulation (Fig. S7-8). As expected, 24h respiratory exchange ratio (RER) profiles were significantly influenced by time of day, reflecting normal circadian variation in substrate utilization, and were significantly impacted by the interaction of TRF x time of day (Fig S7A). While TRF did not have a significant main effect on mean 24h RER, post-hoc analyses revealed significantly lower RER values during the delayed onset of feeding in the 8h-TRF group compared with AL controls in both sexes, consistent with extended lipid oxidation during the prolonged fasting period. Additionally, in males housed at room temperature, post-hoc analyses identified higher nocturnal RER values at discrete time points under 8h-TRF, suggesting a shift toward increased carbohydrate utilization during the night and an enhanced diurnal amplitude of RER.

24h energy expenditure (EE) profiles also exhibited expected diurnal variation and were significantly influenced by TRF in females, with 8h-TRF trending lower during the daytime/rest phase (Fig. S7B). Post-hoc analyses further revealed that in 8h-TRF females housed at thermoneutrality, daytime EE was significantly reduced at several time points, whereas in males, the minimal impact of TRF was largely restricted to light/dark transitions and feeding/fasting periods across both temperature conditions. These results indicate that TRF modulates EE in a time-dependent manner, with effects shaped by sex and ambient temperature. Impacts on EE were further analyzed by ANCOVA to account for body composition (Fig. S8). Homogeneity of regression slopes was confirmed, and while lean mass significantly influenced outcomes in males, TRF effects on EE remained largely time-specific rather than altering overall 24h EE in either sex.

#### Glucose homeostasis is moderately improved by 8h-TRF in males

We measured 12h fasting glucose levels and performed glucose tolerance tests at 6, 12, and 18 months of age at ZT10 to assess glucose homeostasis. Despite being under circadian control and strongly influenced by feeding (*53*), neither 12h-TRF nor 8h-TRF produced consistent reductions in fasting glucose across all ages as seen in our previous study (Fig. S9A) (*9*). Interestingly, 8h-TRF females exhibited lower fasting glucose at 12 months, but at 18 months levels were significantly higher. In mice, unlike humans, fasting glucose typically declines with age, so the higher levels observed at 18 months in 8h-TRF females may in fact reflect a healthier glucose state (*54*). Glucose tolerance tests also revealed sex-specific effects of TRF. 8h-TRF females showed a modest improvement in glucose homeostasis at 12 months of age, but no impacts were found at 6 or 18 months of age in either TRF regimen (Fig. S9, B and C). Males exhibited significant improvements at 6 months in both 12h and 8h-TRF, with modest benefits still apparent at 12 months in the 8h-TRF group. Fasting insulin levels from plasma were not significantly altered by either TRF regimen at any age examined (Fig. S10).

#### Little to no impact on circulating metabolic and inflammatory markers

To determine whether TRF broadly altered systemic metabolic or inflammatory signaling, we also measured circulating levels of leptin, brain derived neurotrophic factor (BDNF), monocyte c chemoattractant protein-1 (MCP-1), tumor necrosis factor α (TNFα), interleukins (IL)-1β, –6, and –10 from 12h fasted plasma collected at ZT10 at 6, 12 and 18 months of age. Overall, TRF had minimal impact on these adipokines, with no sustained differences observed between feeding groups across ages (Fig. S10). An exception was observed at 6 months of age, where males subjected to 8h-TRF exhibited higher plasma BDNF levels than 12h-TRF, suggesting an increased neuroprotective benefit, along with elevated levels of the inflammation-associated chemokine MCP-1. However, these differences were not maintained with aging. Another exception was found at 18 months of age where 12h-TRF males had higher levels of IL-1β and 8h-TRF females had higher levels of MCP-1 suggesting that TRF may have age– and sex-specific effects on inflammatory markers in older animals.

### TRF Slows Age-Related Increases in Frailty Index Values

Frailty, a widely used measure of healthspan in aging mice, integrates 31 physiological and behavioral parameters into a composite index (*55*). Frailty assessments were conducted every 6 months and revealed that both 12h and 8h-TRF significantly reduced frailty index scores at specific ages compared to AL controls (Fig. 4, A-C, and Figs. S11-12). In females, 12h-TRF significantly reduced frailty index scores at 18 months, with hearing loss and body condition assessments defining the key differences between the two groups. 8h-TRF further extended reductions in frailty index from 12 to 24 months, with specific improvements observed in grimace, piloerection, forelimb grip strength, coat condition, ulcerative dermatitis, and body condition scores. Males showed similar trends: 12h-TRF reduced frailty indices between 12 and 18 months, primarily improving grimace and body condition, while 8h-TRF sustained these benefits through 24 months and improved additional parameters, including hearing, piloerection, menace reflex, fur color loss, and coat condition. These results show that TRF significantly lowers age-related frailty in a dose-and sex-dependent manner, with 8h-TRF providing prolonged benefits and improving the greatest number of health measures.

**Figure 4:**
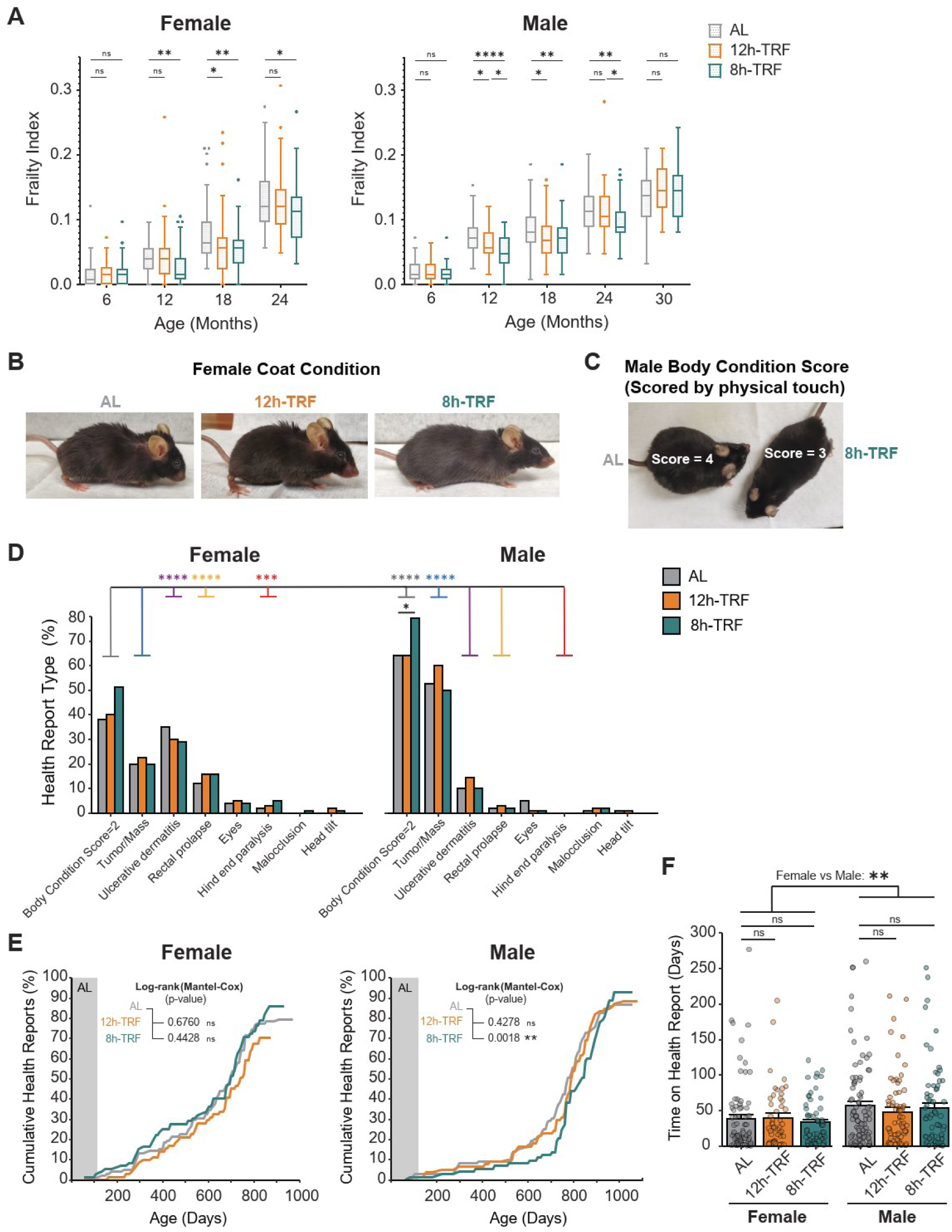
TRF reduces frailty index in both sexes but only delays onset of health reports in males. (**A**) Boxplots of frailty index scores every 6 months. Two-way ANOVA: Females: Age=****, Feeding=****, Interaction=ns; Males: Age=****, Feeding=****, Interaction=**. Asterisks represent significant differences in TRF vs AL (*P ≤ 0.05; **P ≤ 0.01; ***P ≤ 0.001; ****P ≤0.0001) as determined by two-way ANOVA and Tukey’s post-hoc. Females: AL, N=44-79. 12h-TRF, N=34-74. 8h-TRF, N=40-74. Males: AL, N=23-79. 12h-TRF, N=22-78. 8h-TRF, N=39-76. (**B**) One example parameter contributing to reduced frailty index in 8h-TRF females: coat condition, P < 0.05 at 24 months compared to AL. (Full list see Fig. S11). (**C**) One example parameter contributing to reduced frailty index in 8h-TRF males: body condition score, P < 0.05 12-18 months compared to AL. (Full list see Fig. S12). (**D**) Percentage of mice in each feeding group and sex reported to veterinary staff for Body Condition Score=2, Tumor/Mass, Ulcerative dermatitis, Rectal prolapses, Eye ulcerations/swelling, Hind end paralysis, Malocclusions, and Head tilts. Black asterisks represent significant differences in TRF vs AL, and all other colors represent male vs female comparisons as determined by contingency tables and Fisher’s exact tests. (**E**) Cumulative curve showing the percentage of veterinary health reports filed in each group with age. Log-Rank Mantel-Cox Test. (**F**) Time spent on health report until death. Two-way ANOVA: Sex=**, Feeding=ns, Interaction=ns. Data shown as mean ± SEM.

### The Age-Related Onset of Health Reports Reveals Sex-Specific Differences and A Delayed Onset in 8h-TRF Males

As mice developed health issues they were reported to veterinary staff for more frequent monitoring. These reports revealed that TRF did not reduce the relative frequency of commonly reported ailments compared to AL (Fig. 4D). Indeed, 8h-TRF males had significantly more reports of body condition scores declining to 2 (Scored 1-5 with: 1 being emaciated, 2 underconditioned, 3 ideal condition, 4 overconditioned, and 5 obese) (Fisher’s exact test 8h-TRF vs AL: Body Condition Score=2, P < 0.05). However, we observed a clear sex-dependent pattern in the types and frequencies of reported conditions. Females, regardless of feeding group, were significantly more likely to be reported for ulcerative dermatitis (Fisher’s exact test females vs males: P < 0.0001), rectal prolapse (P < 0.0001), and hind limb paralysis (often associated with spinal tumors, P < 0.001). In contrast, males were more frequently reported for external tumors and abdominal masses (Fisher’s exact test females vs males: P < 0.0001), as well as low Body Condition Score=2 (P < 0.0001). The onset of age-related health issues also differed by sex, with 8h-TRF delaying the median health report onset by 52 days in males only compared to AL controls (Log-rank Mantel-Cox, P = 0.0018) (Fig. 4E). Additionally, while there were no significant differences under TRF, overall, females overall spent significantly fewer days on health report prior to death, than males (Fig. 4F). Altogether this demonstrates there is sex-specificity in the development of common husbandry related-disease.

### TRF Has Minimal Impacts on Hematological Markers

To determine whether lifelong TRF impacted hematological markers, which are emerging as predictors of systemic health and aging (*56*), we assessed complete blood counts from whole blood at 6, 12 and 18 months of age. Overall, TRF had minimal effects on circulating blood parameters in either sex. Red blood cell counts (RBC), hemoglobin, hematocrit, red cell indices including mean corpuscular volume (MCV), mean corpuscular hemoglobin (MCH), and mean corpuscular hemoglobin concentration (MCHC), and platelet counts were largely unchanged across feeding regimens (Figs. S13-14). TRF also had no significant impacts on total white blood cell counts or leukocyte subtypes (Fig. S15). The only notable exception was observed in females at 18 months of age, where 8h-TRF mice displayed higher RBC counts accompanied by lower MCV (smaller RBCs; typically indicate older, longer maintained cells) and MCH compared to AL controls (Fig. S13). Given that old age in mice is commonly associated with reduced RBC survival (*57*), this pattern might reflect improved maintenance of RBC homeostasis rather than a pathological shift. Importantly, MCHC, hemoglobin levels, and hematocrit were unchanged in 8h-TRF females, indicating preserved hemoglobin concentration within cells and maintained oxygen-carrying capacity.

### 8h-TRF Extends Lifespan in Males Only

For the first time, we were able to investigate the impact of circadian aligned TRF with regular chow diet, on lifespan in mice (Fig. 5A, and Tables S2-3). In females, no significant extension of median lifespan was observed with either 12h-TRF or 8h-TRF. Median survival was 715 days and 729 days, respectively, compared to 694 days in AL controls. In males, 12h-TRF also failed to extend lifespan, with a median survival of 839 days compared to 818 days in AL, a finding now reproduced across two independent experiments (Fig. S16). Interestingly, 8h-TRF with lifelong self-imposed CR, significantly improved overall survival for males (Log-rank Mantel-Cox, P = 0.0001). Median lifespan increased by approximately 12%, reaching 916 days, and maximal lifespan was prolonged by ∼3% (Fisher’s exact test, P = 0.0038). Though TRF is beneficial for improving health in both sexes, its capacity to extend lifespan appears to be feeding window– and sex-dependent.

**Figure 5:**
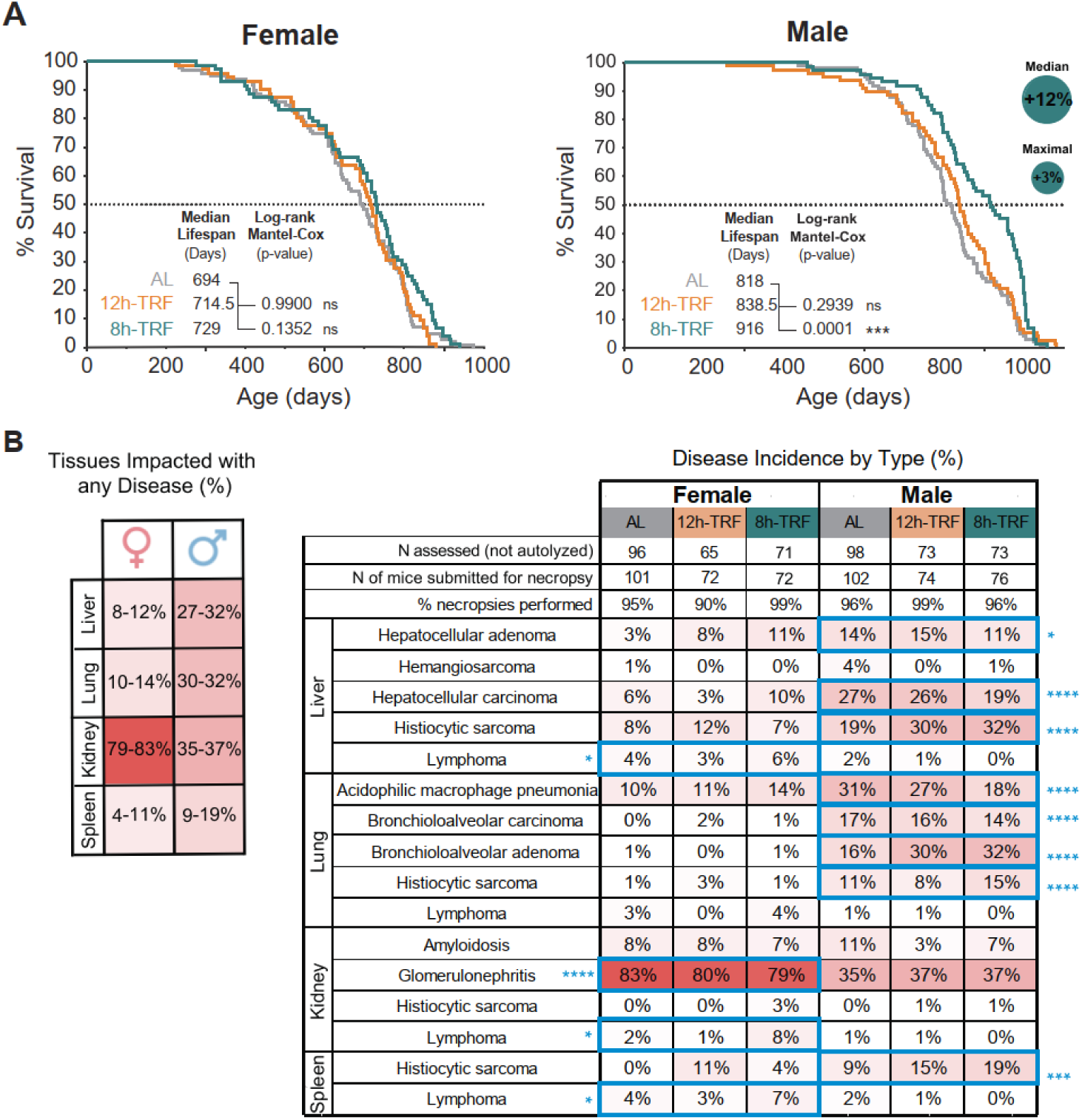
8h-TRF extends lifespan in males. (**A**) Kaplan-Meier survival curves and day median lifespan reached. Log-Rank Mantel-Cox Test for significant difference in overall survival TRF vs AL. Fisher’s exact test for maximal survival TRF vs AL. Females N=72-99 per group. Males N=73-99 per group. (**B**) (*Left*) Histopathology results showing the percentage incidence of any disease type in the most impacted tissue types between females and males and (*Right*) incidence by specific disease type within these tissues separated by sex and feeding group (Full table with all tissues and diseases shown in Tables S4-6). Significant differences in females vs males highlighted by blue boxes and shown with blue asterisks (*P ≤ 0.05; **P ≤ 0.01; ***P ≤ 0.001; ****P ≤0.0001) as determined by contingency tables and Fisher’s exact tests.

### Sex-Dependent Differences in Disease Types and Incidence at Death

After death, each mouse underwent gross necropsy and histopathological analysis by a veterinary pathologist to determine the types of diseases that contributed to its debilitation (Fig. 5B, and Tables S4-6). As shown previously (*10*), the types and relative frequency of diseases were similar for males across all feeding conditions with the most common cause of death or debilitation being neoplasia (Fig. 5B, and Table S7). Along with the delayed onset of health reports, overall, this supports that rather than preventing disease, 8h-TRF delays the development of age-related diseases in males.

However, a few specific neoplastic subtypes were differentially affected by TRF. In males, bronchioloalveolar adenomas, benign tumors, were significantly increased in both 12h-TRF and 8h-TRF groups compared to AL controls. (Fisher’s exact tests AL vs TRF: 12h and 8h, P < 0.05). In females, hepatocellular adenomas, were significantly more frequent in the 8h-TRF group (Fisher’s exact tests AL vs TRF: 8h P < 0.05). While this indicates increased growth of tumors with malignant potential, the absence of a corresponding increase in carcinomas suggests that TRF may shift the spectrum of pathology toward less aggressive neoplastic lesions rather than completely preventing tumorigenesis.

TRF did not reduce the overall occurrence of disease in females either. However, females developed neoplasms far less frequently than males (Fisher’s exact tests females vs males: Adenoma in liver, P < 0.05; Carcinoma in liver and lung, each P < 0.0001; Histiocytic sarcoma in liver and lung, each P < 0.0001; Histiocytic sarcoma in spleen, P < 0.001; Acidophilic macrophage pneumonia in lung, P < 0.0001; Adenoma in lung, P < 0.0001). Instead, most females were debilitated by glomerulonephritis in the kidneys and had a higher incidence of lymphoma than males (Fisher’s exact tests females vs males: Lymphoma in liver, kidney, spleen, each P < 0.05; Glomerulonephritis, P < 0.0001). These results have revealed pronounced differences in neoplasia and development of other age-related morbidities between the sexes.

### TRF Extends Healthspan, With More Prolonged Benefits in Females

While TRF improved and attenuated age-related changes in several markers of health, we next aimed to quantify both the magnitude and duration of its overall impact by measuring healthspan. Frailty index is commonly used to determine healthspan (*55, 58*), but it cannot incorporate the impact of behavioral changes in wheel-activity and food consumption and can miss important physiological changes in the time between assessments. To generate a continuous healthspan index, we integrated frailty with 12 additional health measures by first normalizing each data set (mean = 0, standard deviation = 1), then scaling the parameter directionally by providing context based on whether higher or lower values reflected better health (e.g., high wheel-running activity was scaled positively, while low frailty scores were inverted and also scaled positively) (Fig. 6A), and finally summing the values across age points (Fig. 6B). With higher values reflecting better health, we found that both females and males under TRF had significantly improved healthspan indices throughout life compared to AL controls (Fig. 6, B and C). In 12h-TRF females, this improvement in health was maintained from the onset of TRF until 774 days of age (∼26 months). In 8h-TRF females, health benefits persisted until 879 days of age (∼29 months), and these mice remained healthier than the 12h-TRF group until 858 days (∼29 months). In males, 12h-TRF also improved the healthspan index through 879 days of age (∼29 months), whereas 8h-TRF extended these improvements to 963 days (∼32 months) and were significantly healthier than the 12h-TRF group until 942 days (∼31 months). Importantly, these healthspan indices strongly correlate with lifespan in both sexes, demonstrating that this integrated metric captures biologically meaningful health outcomes (Fig. 6D).

**Figure 6:**
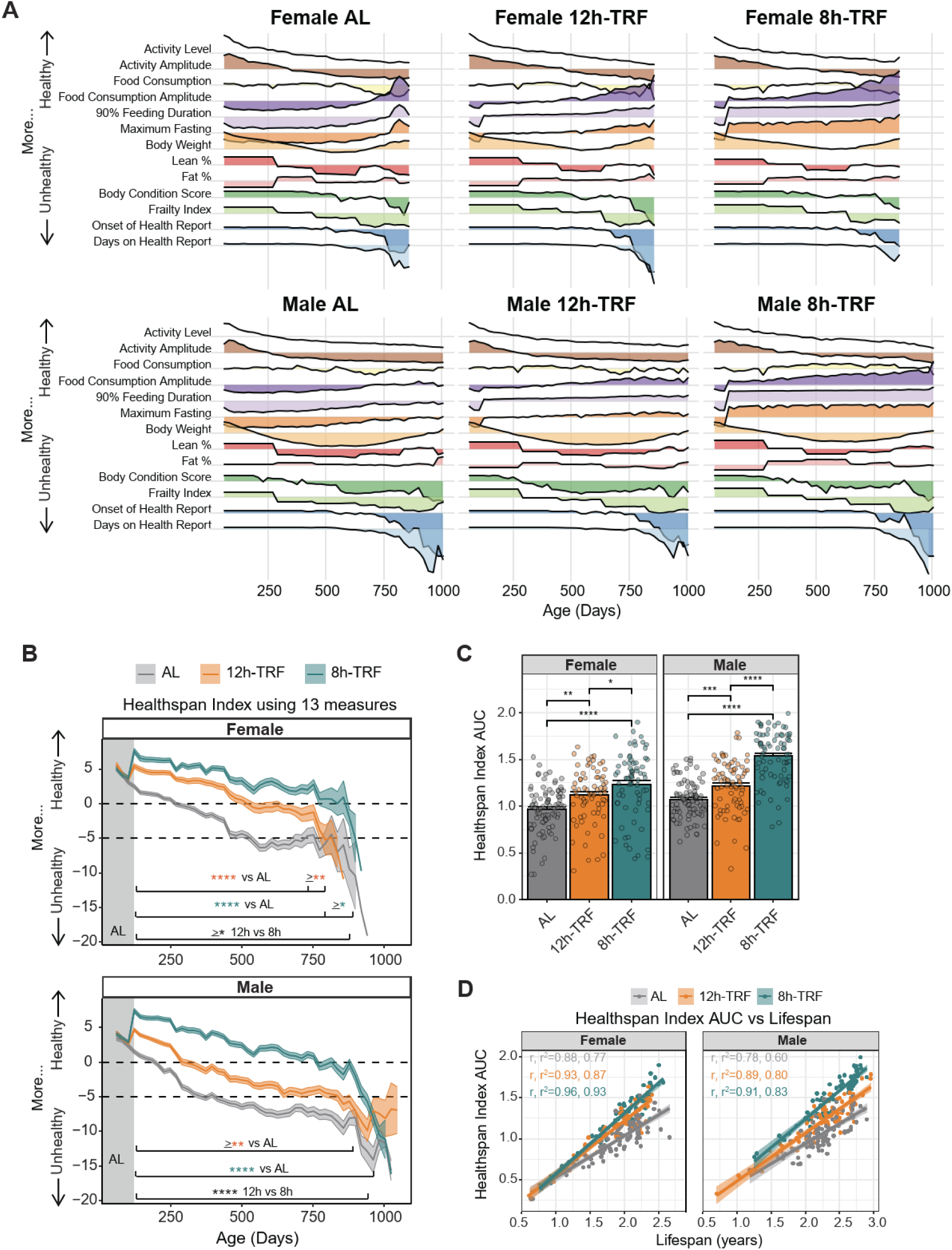
TRF improves healthspan in both sexes. (**A**) Ridgeline plots for each of the 13 health measures recorded throughout the experiment. All parameters were standardized (mean = 0, SD = 1) and, where appropriate, oriented in the positive direction to reflect better health. (**B**) Healthspan index summing the 13 health measures for each group from panel A. Higher values indicate better health and lower values poorer health. Dashed lines (black) mark the age at which each group’s index crosses subjective thresholds of 0 and –5. Type III ANOVA (Wald χ²): Females: Age=****, Feeding=****, Interaction=****; Males: Age=****, Feeding=****, Interaction=ns. Each point represents mean of 21 days ± SEM (shaded regions). Asterisks/percents represent significant differences (*P ≤ 0.05; **P ≤ 0.01; ***P ≤ 0.001; ****P ≤0.0001) as determined by type III ANOVA and Holm’s post-hoc. (**C**) Area under the curve (AUC) for healthspan index in females and males. Type III ANOVA: Sex=****, Feeding=****, Interaction=***. Mean ± SEM (bars). Asterisks represent significant differences as determined by type III ANOVA and Holm’s post-hoc. (**D**) Correlation plots of healthspan AUC vs lifespan in years. Correlation coefficients (r) and coefficients of determination (r^2^) shown.

We also found that health declines were more delayed in females than in males, particularly under TRF. Using subjective thresholds of ‘0’ or ‘-5’ (black dashed lines), AL females crossed the ‘-5’ threshold more than 100 days later than AL males, indicating that females maintain better health for longer, especially relative to their total lifespan. Under 12h-TRF, females not only had prolonged healthspan relative to controls at these thresholds, but also crossed the ‘0’ threshold more than 200 days later than males on the same TRF regimen. While 8h-TRF extended health above the ‘0’ threshold by approximately 500 days relative to AL in both sexes, this also made up a greater portion of the females lifespan. Together, these results demonstrate that TRF extends healthspan in both sexes and further suggest that, despite the absence of lifespan extension in females, females in both TRF regimens remain healthier for a greater proportion of their lifespan than males.

To validate our findings of TRF-induced healthspan extension further, we also applied a previously published summary metric, Frailty-Adjusted Mouse Years (FAMY) (*59*). FAMY, which relies solely on frailty scores, similarly detected a significant healthspan increase in 8h-TRF males (Fig. S17A). The lower frequency of frailty assessments in our study, performed only once every six months, likely contributes to the banded distribution of FAMY scores and may explain discrepancies between the two methods. Nevertheless, FAMY showed strong correlations with lifespan across all groups (Fig. S17B). These results support the robustness of our healthspan measurements and suggest that our more comprehensive, higher-resolution index may be better suited for detecting healthspan changes in females.

## Discussion

Previous studies have established that TRF improves circadian rhythms, ameliorates metabolic syndrome, and improves overall health in obesity models consuming a ‘Western diet’ high in fats and sugars (*28–32*). We expand on these findings by revealing that circadian aligned TRF confers significant, long-term health benefits even under normal nutritional conditions. We show that, depending on the feeding window and sex, TRF prolonged higher amplitudes of daily rhythms in feeding and wheel-running, which are known to feed back on the circadian clock to drive robust molecular and behavioral oscillations (*14, 15, 17, 47, 60–62*). Both TRF paradigms were sufficient to significantly reduce age-related increases in weight and fat mass, having the largest impact during midlife. TRF similarly slowed the age-related decline in lean composition, which may also contribute to the observed improvements in adiposity and suggests that TRF can delay the onset of functional disabilities (*63–65*). We also present novel findings that both TRF windows reduce frailty index in females and males, as in CR and intermittent fasting studies (*66–70*). Veterinary health reports were largely sex-dependent, and overall TRF did not alter the types of deficits observed. Notably, 8h-TRF males displayed a higher proportion of low body condition scores (score of 2), but this occurred alongside a delayed onset of health issues and overall increased lifespan and likely reflects the survival of more mice to advanced ages rather than a detrimental effect of TRF. Consistently, no differences in body condition score were observed at 30 months in males within the individual frailty assessments. Altogether, this demonstrates that TRF does not merely act as a remedial strategy to ‘rescue’ poor health but can actively improve behavioral and physiological markers of health for both sexes under normal metabolic aging conditions.

Despite these robust improvements in health, TRF-induced reductions in body weight and adiposity were not accompanied by consistent improvements in glucose tolerance or sustained alterations in fasting glucose, insulin, leptin, or circulating inflammatory markers. These findings indicate that the metabolic benefits of TRF do not require large shifts in systemic endocrine or inflammatory signaling. Instead, they suggest that TRF may promote health through alternative mechanisms, such as enhanced circadian organization, behavioral rhythmicity, or tissue-specific metabolic adaptations. Consistent with this, 24h EE and RER profiles revealed that TRF primarily reshaped temporal patterns of substrate utilization and EE, rather than altering overall 24h mean values. Importantly, these benefits occurred without detrimental effects on hematological health. TRF had no impact on red blood cell indices, platelet counts, or immune cell abundance across aging, and the modest changes observed in 8h-TRF females were not associated with impaired oxygen-carrying capacity, supporting the long-term physiological safety of circadian aligned TRF.

By integrating measures of frailty, physiology, and behavior into a novel, comprehensive healthspan index, we demonstrate that TRF significantly enhances healthspan in both sexes, with proportionally greater benefits in females relative to their lifespan. We also found that 12h-TRF strongly delayed the initial healthspan decline in females during middle-age, whereas its effect in males at the same threshold is minimal. However, this approach may oversimplify sex-specific differences, as males and females exhibit distinct patterns of health deficits that associate differently with longevity. A single healthspan scale as used here might not adequately reflect these complex, sex-dependent impacts on aging. Although females generally show better health indices, we found they had shorter lifespans than males, highlighting a potential disconnect between this healthspan index and ultimate survival outcomes. It is also important to note, that sex differences observed in mice may not fully translate to humans, as differences in circadian rhythms, hormonal cycles and reproduction, and metabolic rate can alter the magnitude and timing of sex-specific responses to TRF (*71–74*). Regardless, this sex-specific response in healthspan likely reflects a combination of biological factors. Estrogen is known to confer broad protective effects on health and aging (*75*), which may explain the overall female advantage even under AL feeding, and is known to feedback onto the circadian clock (*76–78*). With recent literature showing that females possess a more robust circadian system (*71*), this may underlie their enhanced responsiveness to circadian-based interventions. Recent studies have found that females express a greater number of rhythmic genes, display higher amplitude rhythms, and are more sensitive to external cues (*79–81*). As such, TRF may more effectively enhance rhythms at the molecular level to promote health in females (*19, 20*).

For the first time, we show that aligning feeding to the active phase under 12h-TRF does not extend lifespan. However, a narrower feeding window of 8h-TRF along with self-imposed CR conferred strong lifespan extension in males. These results contrast with previous studies showing that lifelong 20% CR, adjusted relative to age-related changes in AL feeding, extends lifespan by 40.6% in female and 24.4% in male C57BL/6J mice (*82*). While our 8h-TRF mice maintained ∼20% CR for nearly half of their lifespan, both males and females gradually lessened the degree of CR relative to AL feeding shortly before reaching median lifespan. Despite this, 8h-TRF males gained a 12% extension in median lifespan, indicating that even a lower degree of CR is still impactful in males. In contrast, females appear to require a sustained degree of CR ≥20% to see any lifespan benefits. Alternatively, the duration of fasting may be more crucial for female lifespan, as longer fasting periods with CR have enhanced lifespan effects in females (*8, 82*), but not in males (*10*). The mechanisms underlying these sex-dependent differences in response to CR and TRF are not fully understood, especially between different strains of mice (*5, 82*), but are likely influenced by a complex interplay of factors such as differences in nutrient sensing and metabolic pathways, DNA repair, and the effects of sex hormones (*41, 71, 83*).

Survival curves for AL mice unexpectedly showed an 18% increase in median survival for males compared to females, contrary to the common trend in mammalian studies where females typically live longer due to the genetic advantage of a backup X chromosome and the protective effects of estrogen (*84, 85*). However, in C57BL/6J mice, lifespan outcomes are inconsistent possibly due to genetic drift within the isogenic line as well as environmental variations between experiments (*85–88*). While genetic factors may play a role, the shorter lifespan in females could stem from enhanced cold sensitivity and higher thermoregulatory energy demands (*89–92*). The smaller bodied females, needing more energy to maintain body temperature as evidenced by the pronounced increase in food intake with age, may have been adversely affected by the harsher individual housing conditions without nesting material. Although lower body temperatures can extend lifespan (*93, 94*), cold stress can also divert energy away from and impair metabolic, immune, and reproductive functions (*95, 96*). Future studies will be needed to directly evaluate thermogenic pathways underlying these sex-specific effects, including gene expression in brown adipose tissue (*47*). In addition, evaluating TRF and lifespan outcomes under thermoneutral housing conditions or with access to nesting material will help disentangle the relative roles of thermoregulation and feeding timing in shaping healthspan and longevity.

Health report monitoring and post-death histopathological analyses revealed significant differences between the sexes in the progression of age-related diseases, particularly in neoplasia. With the exception of lymphoma, female mice had significantly lower occurrences of neoplasms in the liver, lung, kidney, and spleen which supports a growing body of data that females have greater resistance to cancer (*97–99*). This resistance is largely attributed to sex differences in immunity, mutational burden, and DNA repair, as well as the protective role of X-linked tumor suppressor genes and their interaction with the p53 pathway in females. Although mechanistic experiments are beyond the scope of this study, RNA-sequencing and proteomic analysis of the liver, the tissue most affected by neoplasia in males and least in females, could help to dissect the molecular mechanisms underlying these sex-specific differences in cancer development under TRF. These mechanisms are likely to include pathways regulating the cell cycle and cellular proliferation. Females in this cohort were instead most susceptible to kidney failure due to glomerulonephritis, similar to a previous study (*100*). This condition has been linked to menopause in women, with its progression often accelerating after the decline in estrogen levels (*101*). In mice, this disease can be further exacerbated in the absence of the *estrogen receptor alpha* gene (*102*), emphasizing the protective role of estrogen signaling and uncovering renal disease as a potential biomarker of reproductive aging.

One limitation of this study was the inability of the automated feeders to retract or block a food pellet that goes uneaten beyond the restricted food intake window. This meant that often, a single pellet could be taken by the mouse, outside of its 12h or 8h-TRF window. Additionally, some mice had the tendency to hoard pellets within their cage, leading again to food availability outside of the TRF window, a trait that has already been shown in aged mouse populations (*44*). However, a single daytime pellet accounted for ≤6.34% of total caloric intake, and after subtracting hoarded food, mice consumed ≥90% of the pellets taken from the feeder during the correct feeding windows. These findings, along with evidence that TRF 5 out of 7 days per week regimens still improves metabolic health in mice (*39*), and lifespan in flies (*103*), suggest that moderate TRE adherence may still prolong healthspan and be compatible with modern human lifestyles.

Our study focused on TRF regimens initiated in early adulthood and maintained throughout life, leaving the efficacy of late-life implementation yet to be determined. Short-term studies indicate that TRF in the elderly can reduce high-fat diet-induced inflammation and metabolic dysfunction, and studies with CR in aged mice still provided health and lifespan benefits (*104–106*). However, these benefits provide diminishing returns, dependent on the age at which an intervention is started. Furthermore, as most of the observed benefits occurred through mid-life and the onset of old age, this raises the question of whether a lifelong intervention is necessary, or if stopping TRF before old age might be sufficient, or even preferable, to avoid potential late-life trade-offs. This possibility is supported by other models, where short-term TRF applied in flies earlier in life increases longevity and functional health even after returning to AL feeding (*42, 103, 107*), and by some evidence that TRF in aged mice had negative impacts on lean mass and cardiovascular health (*108*).

Beyond the life stage at which TRF is initiated, the specific timing of the feeding window relative to the internal circadian clock represents a critical variable in metabolic optimization. In this study, the 8h-TRF window was intentionally positioned in the middle of the active period (ZT14–ZT22), instead of the onset (ZT12), as this is consistent with several previous mouse studies (*29, 109*), and prior work in humans suggests that delaying breakfast and/or advancing the final meal can confer metabolic benefits (*110*). Thus, it remains possible that the enhanced healthspan and lifespan benefits observed with 8h-TRF reflect not only feeding duration, but also a shift in the midpoint of daily caloric intake, an aspect of circadian alignment that warrants further investigation. While we observed robust benefits of 8h-TRF in mice, it is also important to note that direct translation to humans is limited by differences in metabolic rate and circadian timing (*73*). In practice, fasting intervals in mice correspond to approximately ∼2.7–3.6 times longer relative windows in humans, so caution is warranted when extrapolating the duration and timing of TRF across species (*73, 74*).

The absence of direct cardiovascular assessments remains a limitation, particularly as cardiovascular disease is a prevalent age-associated pathology in mice. Given the limited data from long-term studies directly evaluating cardiovascular effects of TRF/TRE (*111–113*), future studies incorporating dedicated measures of cardiac function, structure, and blood pressure will be essential to determine how timed feeding influences cardiovascular health across the lifespan, particularly under conditions of normal metabolic aging.

In conclusion, our study provides compelling evidence that the early initiation of circadian aligned TRF either via a 12h or 8h window significantly improves healthspan in both sexes, more so in females proportional to lifespan, and that 8h-TRF robustly extends lifespan in males. While this still warrants further exploration in additional mouse strains and at varying ages of onset, taken together this highlights the potential of TRF as a practical, non-pharmacological intervention to prolong the health quality of human lifespans.

## Acknowledgements

Thanks to G. Martinez, K. Brown, C. Joseph, G. Flowers, A. Torres, A. Herrera, R. Lewis, and L. Thomas, for assisting with the mouse husbandry and experiments. To S. McPherson, K. Goedde, K. Auldridge, and S. Elias Perryman of the Animal Resource Center Diagnostics Lab for their assistance with the gross necropsies. To veterinarians Dr. S. Lewis and Dr. C. Jones for providing guidance on mouse health care. To S. Dixon for providing administrative support. Additional thanks to M. Wellems at Phenome Technologies and D. Ferster at Actimetrics for the feeders and software system. This work was supported by the Howard Hughes Medical Institute awarded to JST, the National Institute of Health (NIH/NIA AG045795 to JST and CBG; NIH/NIA AG072736 to JST and CBG; NIH/NIGMS GM127122 to CBG; NIH 1T32HL138438 to JST and SEI), and the Milky Way Research Foundation MWRF210823 awarded to JST and CBG.

This research was supported in part by the Intramural Research Program of the National Institutes of Health (NIH). The contributions of the NIH author(s) were made as part of their official duties as NIH federal employees, are in compliance with agency policy requirements, and are considered Works of the United States Government. However, the findings and conclusions presented in this paper are those of the author(s) and do not necessarily reflect the views of the NIH or the U.S. Department of Health and Human Services.

## Author Contributions

SEI, NJS, VAR, CBG, and JST conceptualized the study, developed the experimental design, and oversaw all aspects of project execution and supervision. MW-C performed all histopathological assessments and interpretations. SEI, NJS, and VAR conducted all remaining experimental work, including animal handling, data curation, and statistical analyses. SEI, NJS, VAR, CBG, and JST wrote and revised the manuscript with additional edits from MW-C. Funding acquisition was provided by CBG and JST.

## Competing interests

The authors declare no competing interests.

### Data and materials availability

Data are available in the main text or the supplementary materials

## Supplementary Materials

Materials and Methods

Figures S1 to S17

Tables S1 to S7

## Materials and Methods

### Animals and Housing Conditions

For a minimum detection of 10% life extension with 80% power a minimum of 96 animals were needed in the control group and 72 in each of the two TRF groups (*114*). Extra mice were included in each group to account for any that may not adapt to the feeder/diet or for non-aging-related deaths.

264 male and 264 female C57BL/6J mice 6 weeks of age were ordered from Jackson Laboratories, Bar Harbor, ME. We selected C57BL/6J mice because they are widely used research model which exhibit robust circadian rhythms and well-known aging phenotypes (*115–118*). Further, using a single, well-characterized strain maximized statistical power and lifespan resolution in our large cohort (*114*). As done previously, starting at 2 months of age the mice were: **(i)** individually housed in standard polycarbonate cages with stainless steel running wheels inside isolation cabinets under light:dark (LD) of 12:12 hours and ambient building temperature 72-78°F, **(ii)** fed 300mg pellets of purified diet (F0075, Bio-Serv) using automated feeders with water provided *ad libitum* (AL), and **(iii)** cage changed every 21 days (*9, 10, 119*). Nesting material and igloos were excluded, besides shavings, to prevent wheel blockages. The Institutional Animal Care and Use Committee (IACUC) of the University of Texas Southwestern Medical Center approved the animal protocol (APN 2015-100925), which has been renewed every 3 years (in 2018, 2021, and 2024).

The purified diet (F0075, Bio-Serv) is composed of 18.7% protein, 5.6% fat, 4.7% fiber, and 59.1% carbohydrates, similar to regular mouse chows. Major ingredients include Sucrose. Dextrose, Casein, Corn Oil, Mineral Mix, Cellulose, Corn Syrup, Calcium Silicate, Vitamin Mix, Magnesium Stearate, Choline Bitartrate, DL-Methionine, L-Cystine, Ascorbic Acid, Vitamin E Acetate, and tBHQ.

An additional cohort of C57BL/6J animals (N=48/feeding condition/sex) were included in this study in order to undergo more invasive assays. These mice were randomly selected (N=8-12/group/age point) for analysis of metabolism in metabolic chambers, fasting blood glucose levels, glucose tolerance, biomarkers of inflammation and metabolism and hematology. These mice were not included in the longitudinal health measurements or the assessment of lifespan.

### Time-Restricted Feeding (TRF) Regimens

At 2 months of age, unmated mice were placed in individual cages as described above and fed *ad libitum* (AL) for 8 weeks using the automated feeder system (*9, 10*). At 4 months of age, the mice were divided into three feeding groups for the remainder of the experiment: **(i)** 108 of each sex continued in AL as a control group, **(ii)** 78 of each sex in a 12-hour TRF group where the automated feeder restricted food dispensing to the 12-hour night (12h-TRF, ZT12-24), and **(iii)** 78 of each sex in an 8-hour TRF group with food dispensing restricted to the middle 8 hours of the night (8h-TRF, ZT14-22). These feeding windows were chosen as they have been previously validated and demonstrated to confer health benefits in mice (*25, 29*). In addition to the timing of the feeding window, the feeders were also programmed with a 10-minute delay between pellet dispensing to reduce hoarding behavior. Regardless of the feeding group, all mice had access to a maximum of 22 pellets/day or 6.6 g of food. In this study, no group exceeded 6.6 g daily consumption. Previous studies have shown that C57BL/6J mice consume 4-5g of food per day (*120*), so this amount of food was chosen to ensure mice were not calorically restricted.

### Exclusion of Hoarded Food

Mice exhibit a natural tendency to hoard food within the cage, especially later in life or when ill (*44*). In all TRF groups, mice identified as hoarders were flagged, and hoarded food was removed from the cage on a weekly basis. For mice on veterinary health report, hoarded food was removed at their twice weekly full health check. The total amount of hoarded food was quantified by weight (g) at each 21-day cage change. For AL mice, hoarded food was allowed to remain in the cage until the scheduled cage change, at which point it was collected and quantified. In all groups, the total hoarded food weight was divided by 21 days to estimate an average daily hoarding amount, which was then subtracted from the average daily food retrieval recorded by the feeder.

### Daily Monitoring of Feeding and Wheel-Running Activity

As before, the feeders were set and continuously recorded quantity and timing of food intake via ClockLab Chamber Control Software v3.401 (Actimetrics Inc., Wilmette, IL, USA) (*9, 10*). The feeding data presented in Figure 1B and Supplemental figures S2 and S3 represent food consumption with hoarding removed. All other feeding figures represent the quantity and pattern of pellet taking from the feeder, not excluding hoarded food. Wheel-running activity was also continuously recorded as before using an updated version of ClockLab Data Acquisition System v3.604 (Actimetrics Inc., Wilmette, IL, USA).

### Body Weight and Composition Measurements

The body weight of each mouse was measured every 21 days during cage change (morning, ZT4-7) throughout its lifespan. Body composition measurements of lean and fat mass (g) were assessed every 6 months during cage change (ZT4-7) using a 100H-EchoMRI Body Composition Analyzer (EchoMRI, Houston, TX, USA).

### Promethion Metabolic Chambers

These tests were performed in a follow-up cohort of mice. Mice were randomly selected.

At 19 months of age, mice received body composition measurements (EchoMRI) and then were individually housed in metabolic cages (Promethion, Sable Systems International) modified with automated feeders to maintain the same feeding regimens. These cages were housed in light-tight, temperature-controlled cabinets. Mice were acclimatized for 2-3 days at room temperature (25°C) before recording for an additional 5-6 days at room temperature. Cabinet temperature was then raised to thermoneutrality (30°C) for 5 days.

### Fasting Glucose and Glucose Tolerance Testing

These tests were performed in a follow-up cohort of mice. Mice were randomly selected from the cohort at each age point.

For fasting glucose, cages for all mice to be tested were changed out fresh the day before. Feeders for all groups were shut off at ZT21.5 to allow the mice 30 minutes to consume the last pellet dropped and officially begin their 12h fast at ZT22. Mice were checked again at ZT0 to ensure no pellets remained in the cage or available in the pellet chute. At ZT10, a drop of blood was measured from the tail using an Accu-Check Glucose Meter.

For glucose tolerance testing, cages were changed fresh and mice fasted as described above. At ZT10, mice were injected with a 20% glucose solution at a dose of 0.1mL per 25g of body weight. A drop of blood from the tail was measured for glucose at the time of injection (0 min) and then at 15-, 30-, 45-, 60-, and 120-min post-injection.

### Biomarkers of Metabolism and Inflammation

These markers were measured in a follow-up cohort of mice. Mice were randomly selected from the cohort at each age point.

Mice were switched to fresh cages and were fasted for 12h as described above for fasting glucose. At ZT10, 70-100µL of blood was collected via submandibular bleed into K_2_ EDTA coated collection tubes (RAM Scientific Safe-T-Fill™ Capillary Blood Collection Systems) with cOmplete™, Mini, EDTA-free Protease Inhibitor Cocktail (Roche) and DPP-IV (Millipore).

Blood was immediately kept on ice and then spun down at 2000g for 10 min at 4°C. The upper layer of plasma was transferred to a new tube and stored at –80°C until use.

20-25µL of plasma was then measured for leptin, insulin, IL-1β, IL-6, IL-10, TNFα, MCP-1, and BDNF using a U-PLEX Adipokine Combo 1 (mouse) assay kit (Meso Scale Discovery).

### Hematology

Measurements for red blood cells, white blood cells, and platelets were performed in a follow-up cohort of mice. Mice were randomly selected from the cohort at each age point.

Mice were switched to fresh cages and were fasted for 12h as described above for fasting glucose. At ZT10, 25-30µL of blood was collected via submandibular bleed into K_2_ EDTA coated collection tubes (RAM Scientific Safe-T-Fill™ Capillary Blood Collection Systems) with cOmplete™, Mini, EDTA-free Protease Inhibitor Cocktail (Roche) and DPP-IV (Millipore). 20µL of whole blood was then measured in a Hemavet 950FS (Drew Scientific) at room temperature within 2-4 hours of collection.

### Frailty Scoring

Every 6 months the mice were assessed for frailty, or physical vulnerabilities, according to 31 parameters of aging modified from Whitehead et al. 2014 (*55*). These parameters include a range of physiological evaluations, scoring coat condition, ocular and auditory responses, physical and musculoskeletal state, respiration rate, body weight, and temperature. Each assessment was scored either 0=absent, 0.5=mild, or 1=severe. For scoring Body Condition Scores, 0.5 was used for either a BCS of 2 or 4, and 1 was for a BCS of 1 or 5. For body weight and body temperature scores only, these were determined by averaging the group measurements and then determining how many standard deviations (SD) each individual mouse was from their age-, sex– and feeding-matched group mean. If <1 SD, mouse scored as 0 for that parameter, 1-1.99 SD = 0.25, 2-2.99 SD = 0.5, 3-3.99 = 0.75, and >4 SD = 1. The scores were totaled to calculate an overall index score.

### Health Monitoring and Survival Study

Mice were physically health checked every 21 days during cage change and visually inspected every 10 days during refilling of the automated feeders. Using the feeding and wheel monitoring software, mice were also checked virtually every day and only physically health checked if feeding fell to <5 pellets per/day and/or wheel running fell below set thresholds (1.14 counts/min within 24hr, reduced by 10% every 6 months as the mice age) as performed previously (*10*).

If any mouse was found to have a mild/moderate health ailment (e.g. dermatitis, external tumor, abdominal mass, head tilt) or a body condition score < 3 (Scored 1-5 with: 1 being emaciated, 3 ideal condition, and 5 obese) it was placed on veterinary health report. These records were also used to generate Figures 4D-F. Mice on health report were physically health checked weekly by the UT Southwestern Animal Resource Facility (ARC) veterinary staff and twice a week by our own laboratory staff. In order to obtain accurate but humane lifespan data, mice were monitored carefully for euthanasia criteria that would indicate imminent death (moribund) or unacceptable levels of pain (analgesics were not used to avoid confounding lifespan results). Criteria were determined in conjunction with the ARC veterinarians according to AAALAC guidelines and include: Non-responsive to touch, hypothermic or cold to touch, slow or labored breathing, failure or inability to eat/drink after moist chow is given, a body condition score of 1, body weight loss of >20% from baseline or between cage changes, a broken/fractured/dislocated limb, head tilt that is resulting in an animal being unable to maintain sternal body position which would affect their ability to consume food and water, severe eye protrusion/infection, severe dermatitis, grade 3 rectal or penile prolapse, urinary obstruction, or a subcutaneous tumor >2cm.

The number and date of those euthanized or found dead across the feeding groups and sexes were recorded and used to generate the Kaplan-Meier survival curves. A total of 35 animals (35/528) were censored from the survival curve if death occurred earlier than 6 months or due to non-aging related injuries or death: 9 AL females, 6 12h-TRF females, 6 8h-TRF females, 9 AL males and 5 8h-TRF males.

### Necropsy and Histopathology

Mice euthanized or found dead were submitted for gross necropsy and blinded histopathological analysis was performed, as done previously (*10*). Less than 10% of the animals submitted were too autolyzed to analyze. Disease types were tallied together by tissue type, sex, and feeding group.

### Healthspan Index

Throughout the experiment we recorded thirteen health measures including: wheel-running activity to determine overall activity level and diurnal amplitude, food consumption to determine overall level and diurnal amplitude, 90% feeding duration, maximum fasting times, body weight, composition of lean and fat mass as a percentage of body weight, body condition score, frailty index, and the onset and duration of veterinary health reports.

Healthspan index was calculated by summing the calculated scaled values of each of the thirteen health measures. In the case of body condition score, an absolute difference from BCS=3 was calculated. For body fat percentage, an absolute difference in value from 24.45% was calculated. The value 24.45% was used based on previous literature showing the average body fat percentage across multiple healthy mouse strains at the age of 16 weeks (*121*). All measures were then normalized to a mean of 0 and a standard deviation of 1, within each measure. Certain values were then inverted, given their context, in order to have positive numbers representing a healthier state, and negative values representing an unhealthy state. For example, “Days on Health Report” is inverted, as higher values would be less healthy, and lower values are healthier. All normalized values were then summed together at 21-day intervals.

### FAMY

FAMY was calculated using the lifetime of frailty scores collected for each mouse and as described in Lamming 2024 (*59*).

### Statistical Analyses

All measurements represent repeated measures from the same animals over time, with the exception of data shown in Figs. S7–S10 and S13–S15, which were collected from a separate cohort using different mice at each age point. Sample sizes (N) for each experiment are indicated in the corresponding figure legends.

As performed previously, all feeding and wheel-running activity was collected using Chamber Control and ClockLab, respectively (Actimetrics, Inc) (*10*). Each point on the longitudinal profiles represents a 21-day mean. 24h profiles were binned by hour, with each age point reflecting a 14-day mean (taken from the middle of each 21-day cage cycle to avoid behavioral changes due to cage change stress). For each mouse, each 14-day bin was then analyzed by 1-dimensional Discrete Fast Fourier Transform (FFT) using the mixed-radix algorithm for real-valued input (SciPy v1.12.0; scipy.fft.rfft) (*122*). The Blackman window function is applied to the data prior to FFT analysis to inhibit spectral leakage (*123*). The power of each frequency component is then normalized, with the total normalized power of all frequency components equaling 100. The normalized power from the 24h frequency component is then taken as the diurnal amplitude of either food intake behavior or running wheel activity. This value represents the percentage of contribution of the 24h frequency component, to the entire 14-day behavioral period. For example, if the diurnal amplitude for food intake is 21, it indicates that 21% of the observed variation in food intake can be directly attributed to the animal’s consistent 24-hour daily cycle. Group-wise statistics were then performed to determine the overarching impact of each intervention, as described below.

For the calculation of the 90% feeding window, food intake data was similarly partitioned into 3-week intervals, with each sampling point consisting of a 14-day window. Only windows containing a minimum of 84 feeding events (averaging 6 events per day, or ∼1.8g of intake) were included in the final density analysis to ensure robust statistical estimation. Because time-of-day is a periodic variable, we employed a von Mises distribution (the circular analogue of the Normal distribution) to estimate the probability density of feeding activity. We utilized the vonmises.pdf method (SciPy v1.12.0), to compute the circular distribution, where the concentration parameter κ, was defined by a fixed bandwidth of 0.04363 radians, representing approximately 10 minutes of smoothing. To define the primary feeding window, we identified the shortest continuous arc on the circle containing 90% of the total probability mass. This method accounts for feeding windows that cross over the ZT24/ZT0 boundary.

Cumulative health report and survival curves were analyzed using Log-rank Mantel Cox and Fisher’s exact test, health report type and necropsy report disease type percentages analyzed by contingency tables and Fisher’s exact test and, and all other plots analyzed using two-way ANOVA or type III ANOVA with Tukey’s or Holm’s post-hoc analyses (*124*). All statistical tests were two-sided. Normality was assumed but not formally tested. Data plots were generated and statistically analyzed by Prism 10 (Graphpad Software Inc), Python, and R software, using custom-written scripts and standard libraries.

## Supplemental Figures and Legends

**Figure S1:**
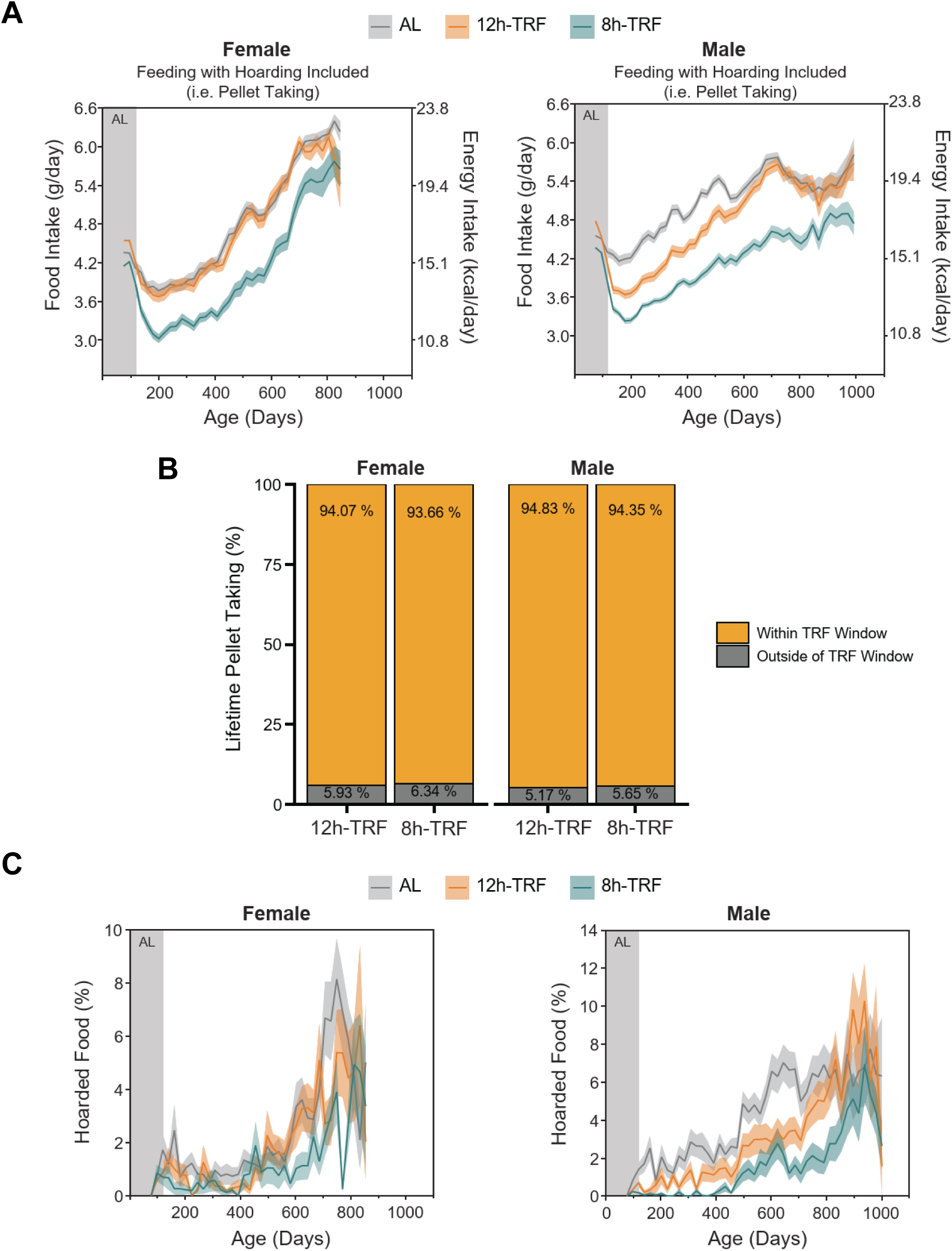
Food taking and hoarding behavior: 300mg pellets are dispensed from the feeder one at a time for the mouse to retrieve. In females and males: (**A**) Longitudinal profiles of daily food taken from the feeder including uneaten (hoarded) food found in the cage (g/day) and the equivalent energy intake (kcal/day). (**B**) Lifetime averages of the percent (%) of food taken from the feeder within or outside of the TRF window. (**C**) Longitudinal profiles of hoarded food removed from cages as a % of the total food taken from the feeder. <u>A and C:</u> *Ad libitum* (AL) baseline feeding (Gray area). Time-restricted feeding (TRF) (White area). Mean of 21 days ± SEM (shaded regions). Females N=71-103 per group. Males N=71-97 per group.

**Figure S2:**
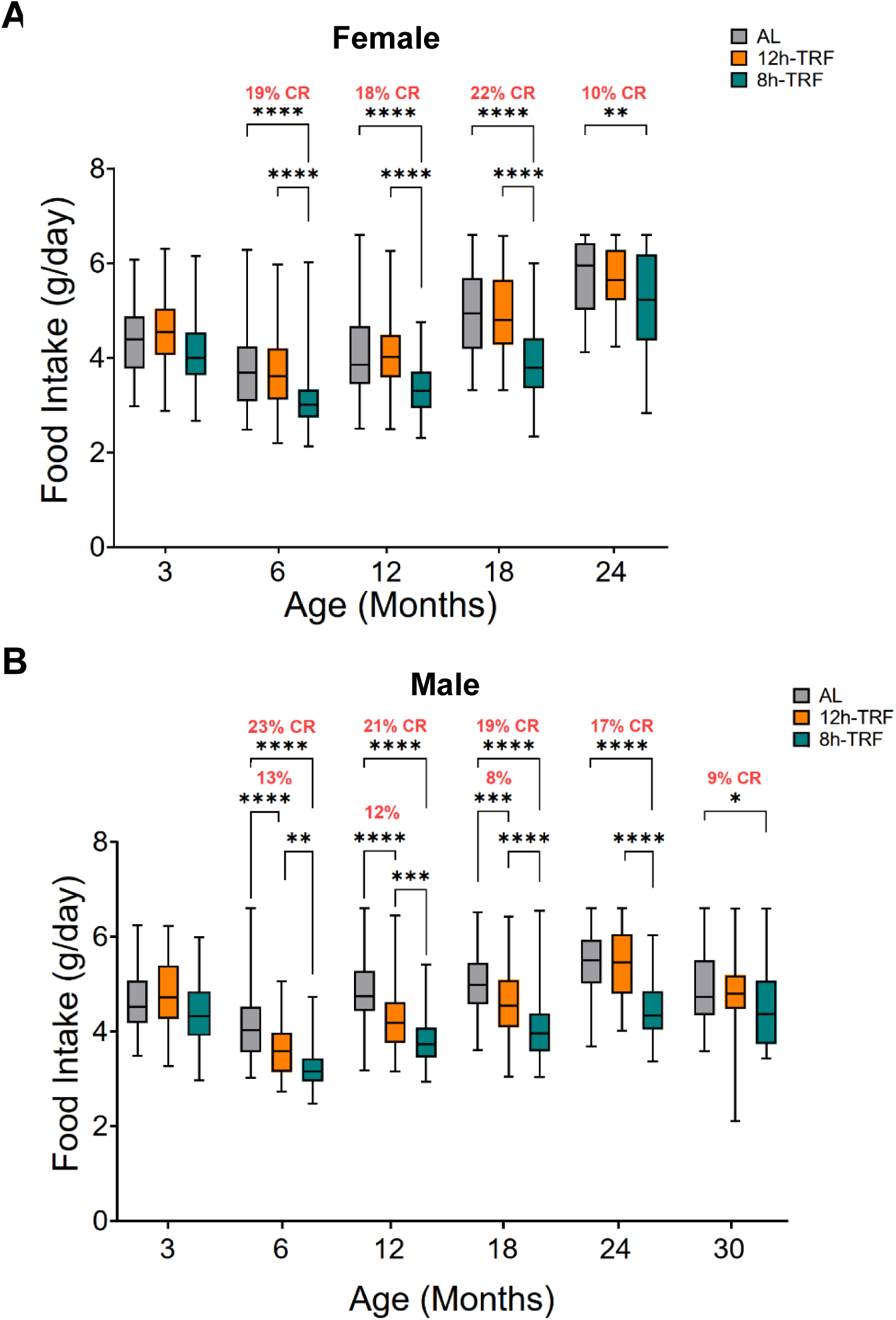
Food intake by monthly age epochs: (**A**) In females and (**B**) males, box plots of food intake with hoarding removed (g/day) at 3 and 6 months of age and then every 6 months following. Food intake per mouse averaged every 21 days. Any caloric restriction (CR) relative to the AL control at each age epoch shown as %. Two-way ANOVA, Tukey’s post-hoc. *P ≤ 0.05; **P ≤ 0.01; ***P ≤ 0.001; ****P ≤0.0001.

**Figure S3:**
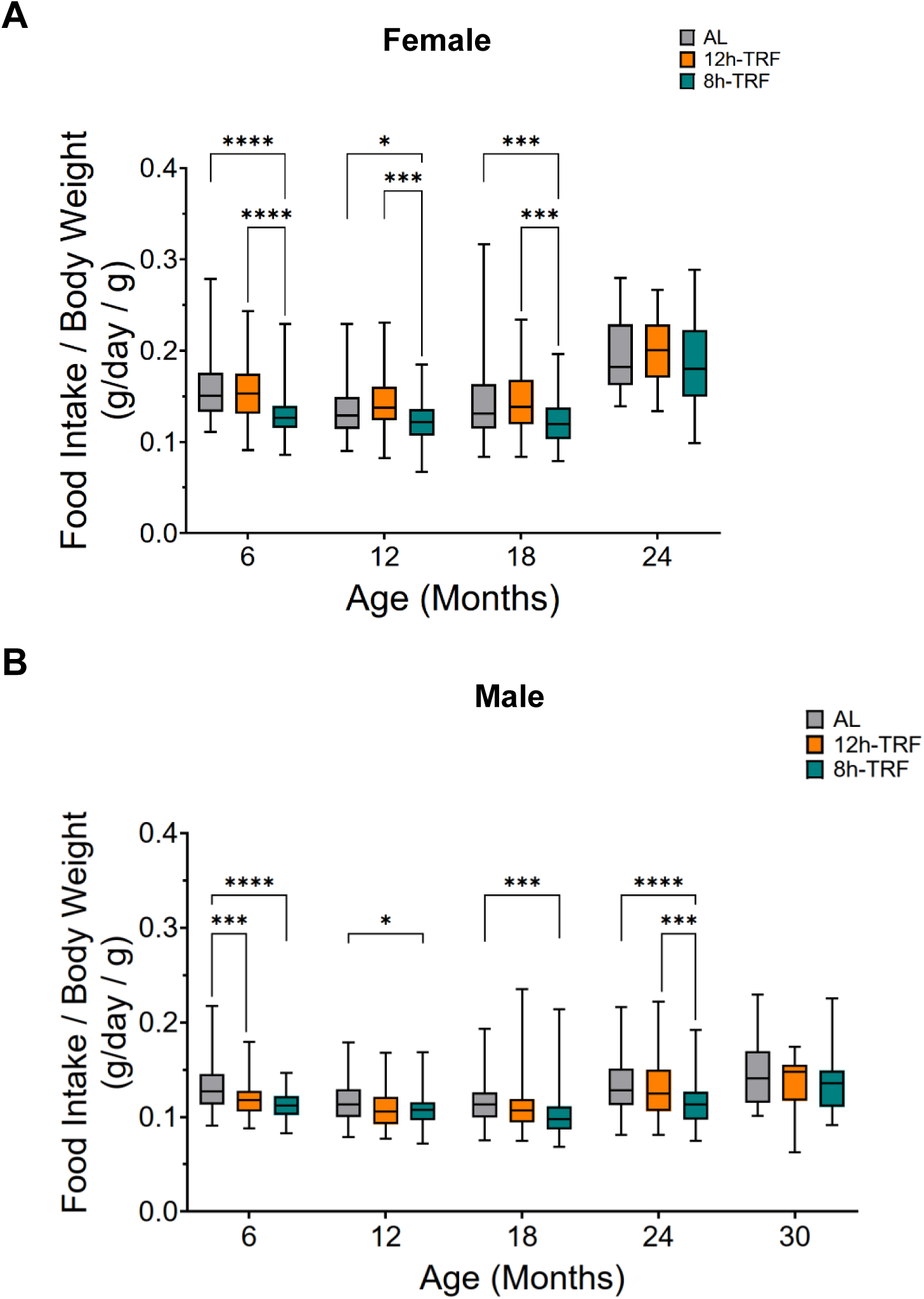
Food intake relative to body weight by monthly age epochs: (**A**) In females and (**B**) males, boxplots of food intake (g/day) with hoarding removed relative to body weight (g) at 6 months of age and then every 6 months following. Two-way ANOVA, Tukey’s post-hoc. *P ≤ 0.05; **P ≤ 0.01; ***P ≤ 0.001; ****P ≤0.0001.

**Figure S4:**
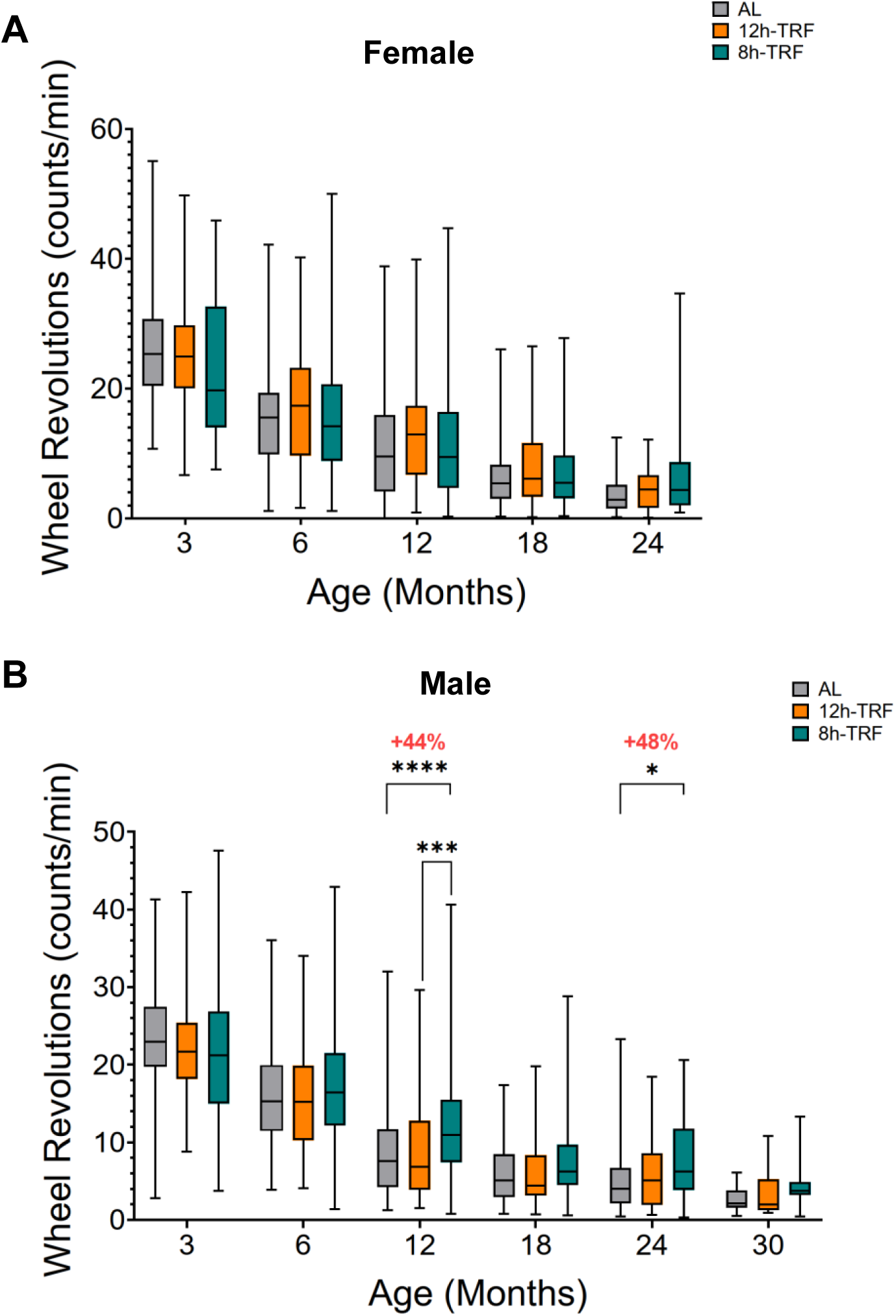
Wheel activity by monthly age epochs: (**A**) In females and (**B**) males, boxplots of daily wheel revolutions (counts/min) at 3 and 6 months of age and then every 6 months following. Wheel running per mouse averaged across 21 days at each age point. Two-way ANOVA, Tukey’s post-hoc. *P ≤ 0.05; **P ≤ 0.01; ***P ≤ 0.001; ****P ≤0.0001.

**Figure S5:**
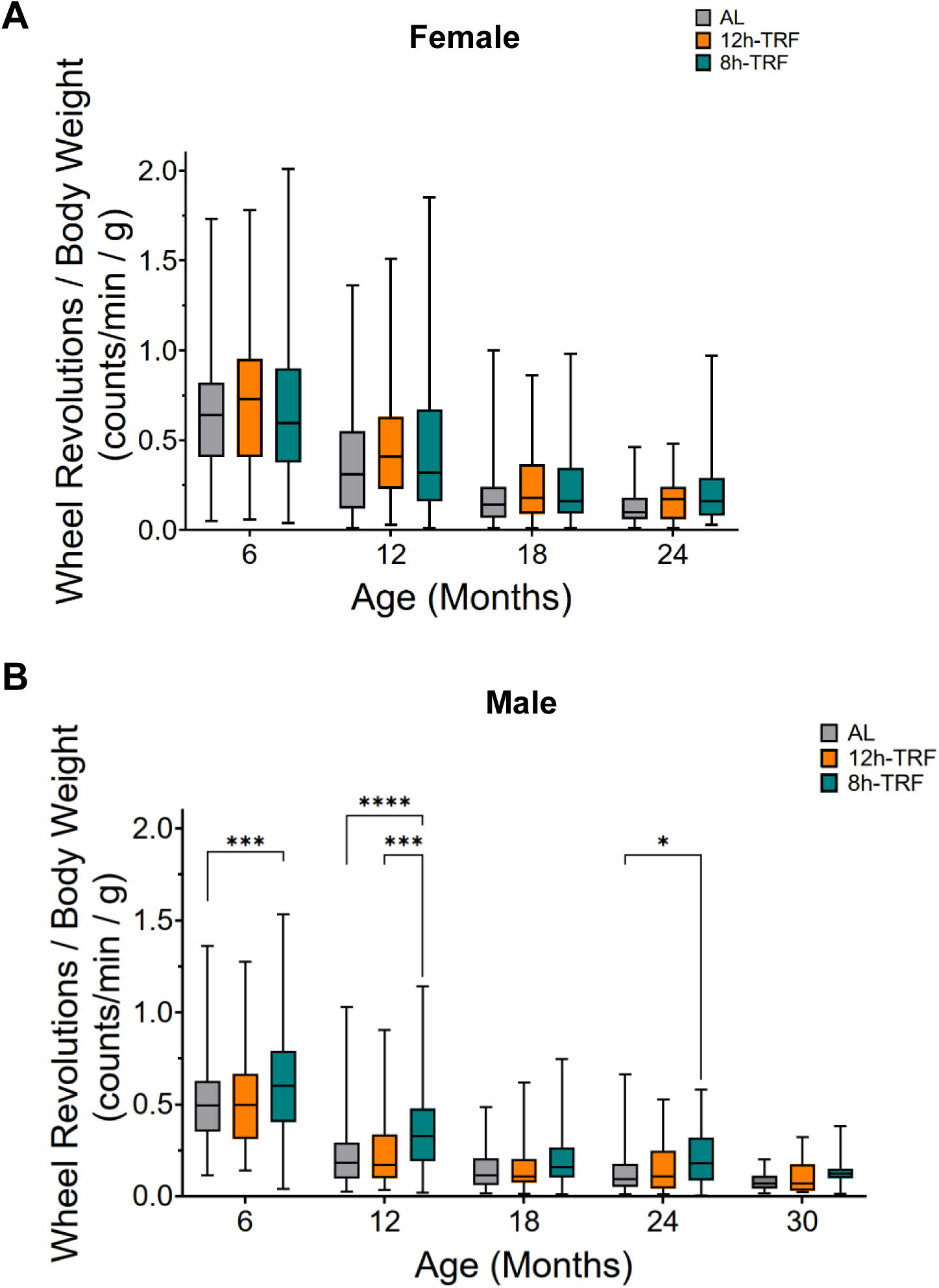
Wheel activity relative to body weight by monthly age epochs: (**A**) In females and (B) males, boxplots of wheel revolutions (counts/min) relative to body weight (g) at 6 months of age and then every 6 months following. Two-way ANOVA, Tukey’s post-hoc. *P ≤ 0.05; **P ≤ 0.01; ***P ≤ 0.001; ****P ≤0.0001.

**Figure S6:**
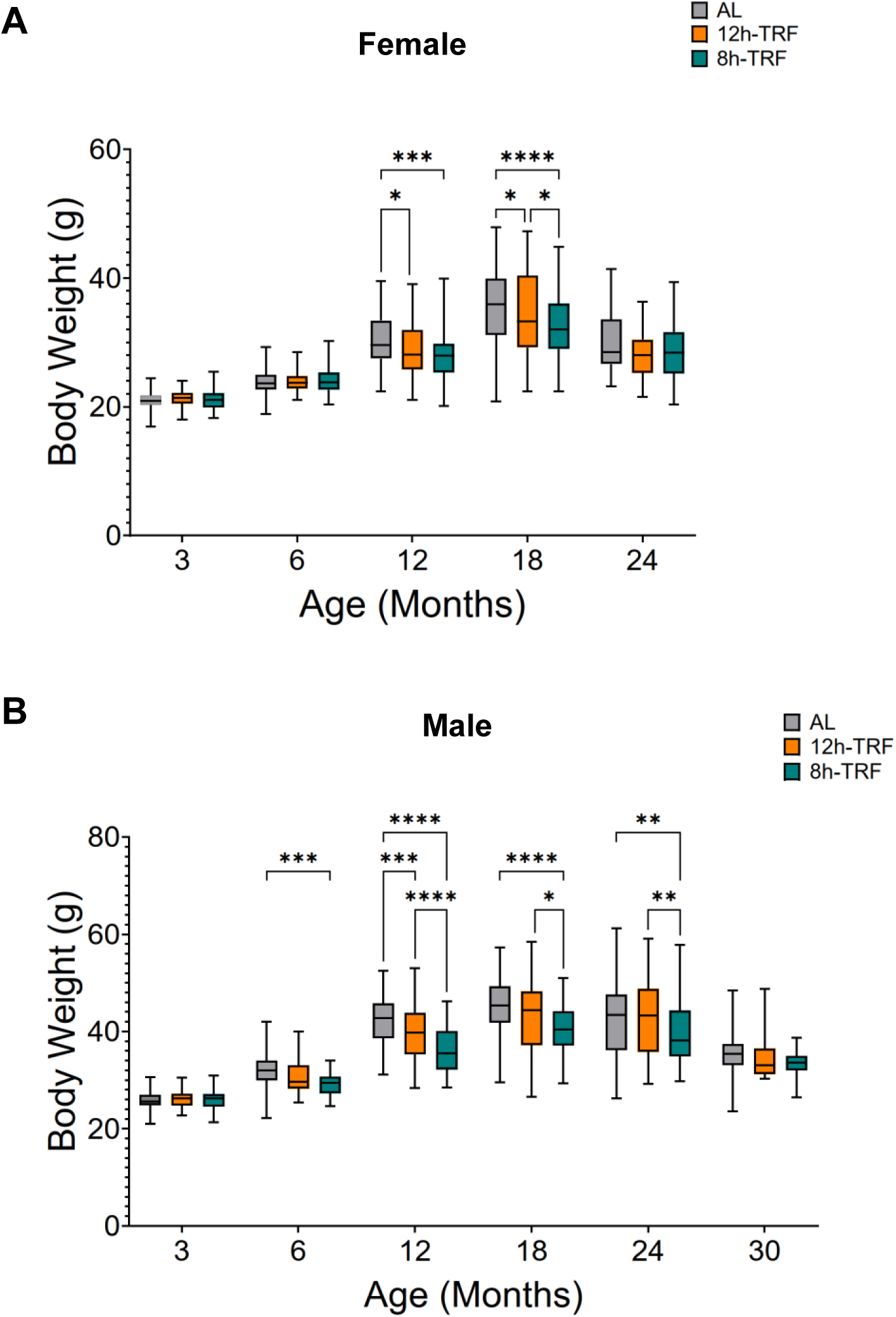
Body weight by monthly age epochs: (**A**) In females and (**B**) males, boxplots of body weight (g) at 3 and 6 months of age and then every 6 months following. Two-way ANOVA, Tukey’s post-hoc. *P ≤ 0.05; **P ≤ 0.01; ***P ≤ 0.001; ****P ≤0.0001.

**Figure S7:**
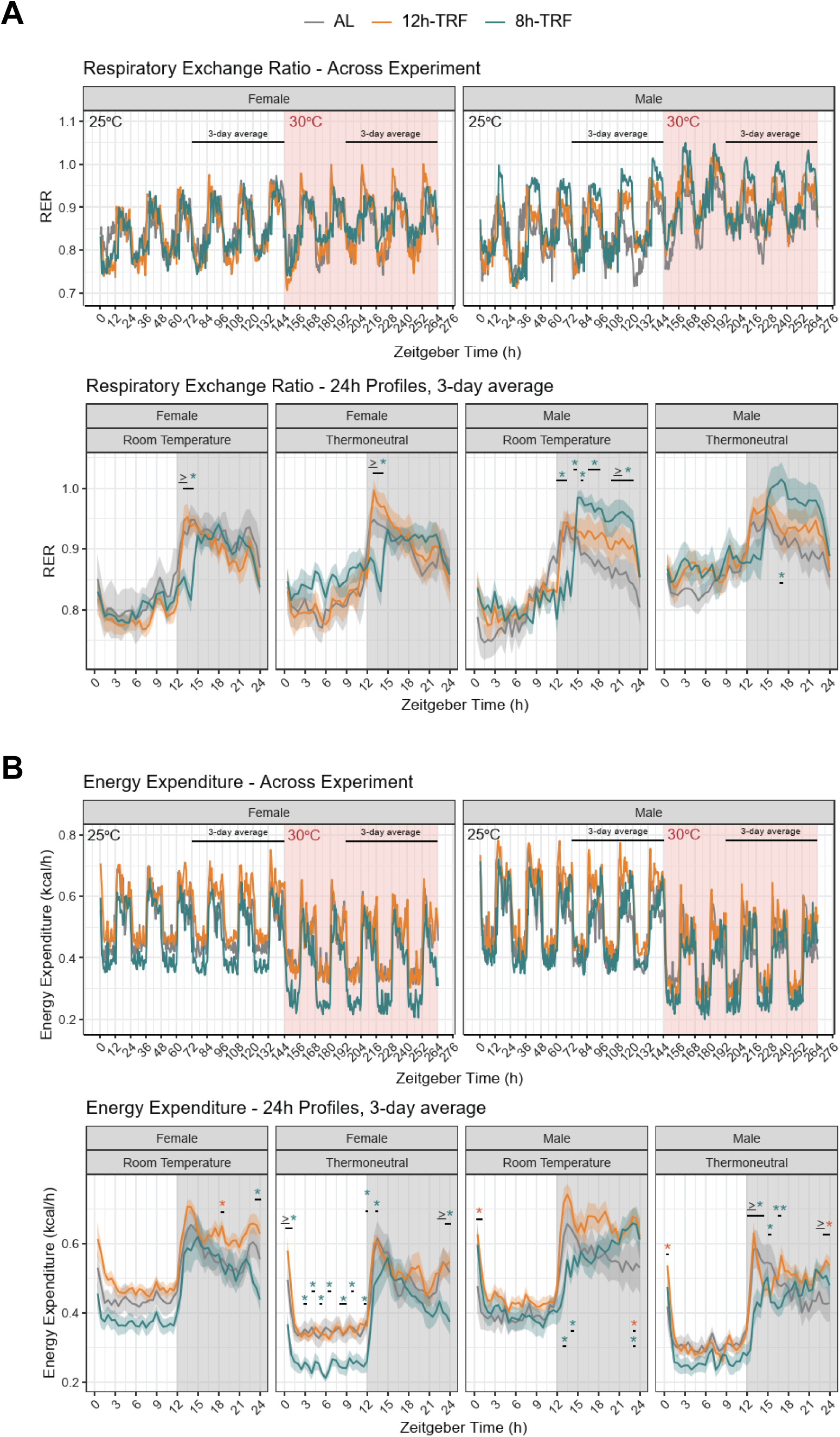
Measures of metabolism at 19 months of age in a follow-up cohort of mice: (**A**) (*Top*) Respiratory exchange ratios (RER; VCO2/VO2) in females and males throughout a 11-day recording in Promethion metabolic chambers. 6 days recorded at room temperature (25°C; white shaded area), and 5 days were recorded at thermoneutrality (30°C; red shaded area). (*Bottom*) 24h profiles of RER averaged from the last 3 days of recording in each temperature. 12h light (white shaded area), 12h dark (gray shaded area). Means ± SEM (shading) presented in 30-minute bins. (**B**) (*Top*) Energy expenditure (EE; kcal/h) in females and males throughout the 11-day recording. 6 days recorded at room temperature (25°C; white shaded area), and 5 days were recorded at thermoneutrality (30°C; red shaded area). Measurements are presented in 30-minute bins. (*Bottom*) 24h profiles of EE from the last 3 days of recording in each temperature. 12h light (white shaded area), 12h dark (gray shaded area). Means ± SEM (shading) presented in 30-minute bins. <u>A and B:</u> Type III ANOVA (Wald χ²), and Holm’s post-hoc. *P ≤ 0.05; **P ≤ 0.01; ***P ≤ 0.001; ****P ≤0.0001.

**Figure S8:**
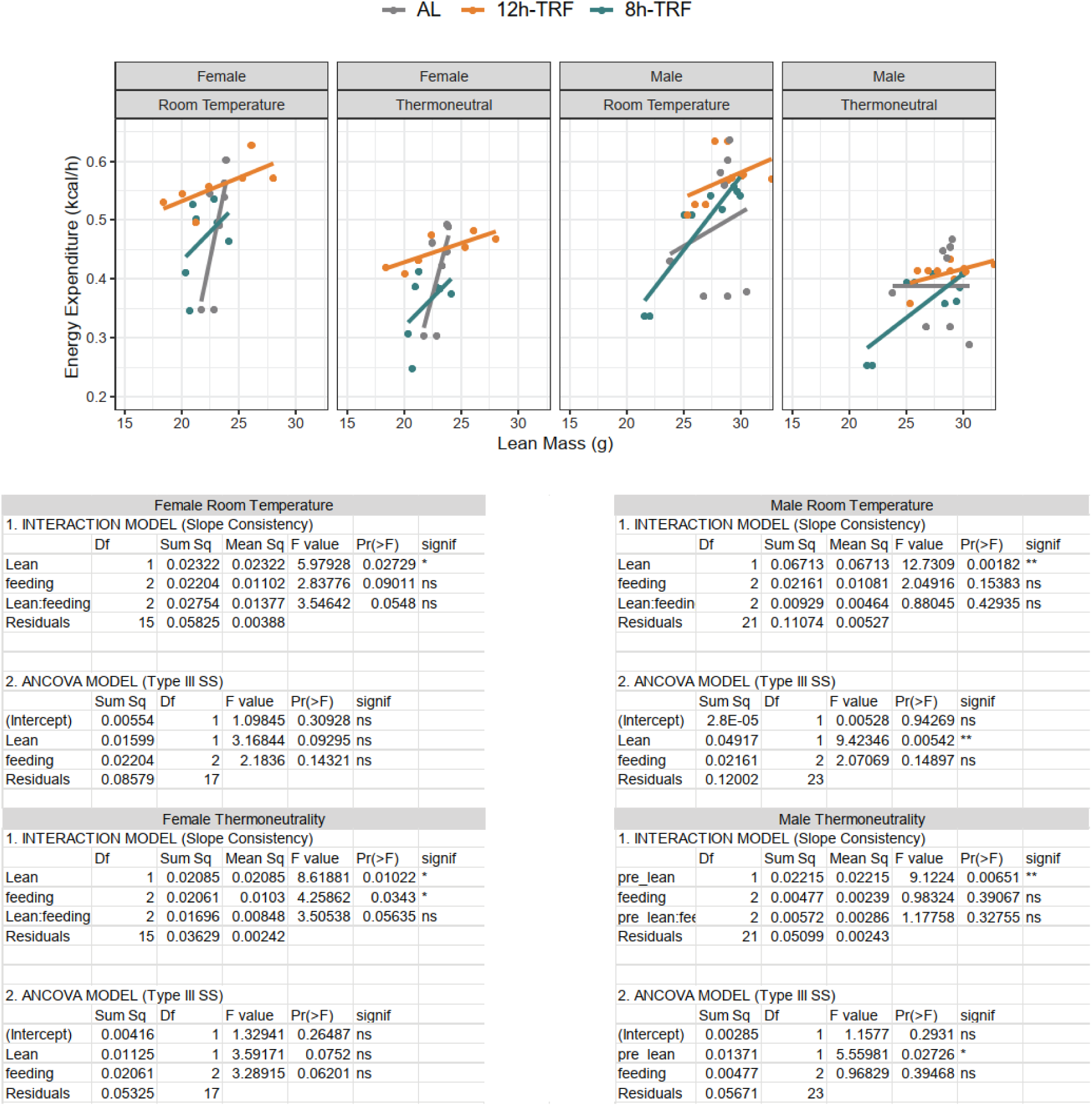
ANCOVA of TRF effects on energy expenditure: 24h mean EE plotted vs lean mass for females and males. Homogeneity of regression slopes was confirmed for all groups (test of Feeding × Lean Mass interaction, all p > 0.05, ns), validating the use of ANCOVA to test for significance of intercepts.

**Figure S9:**
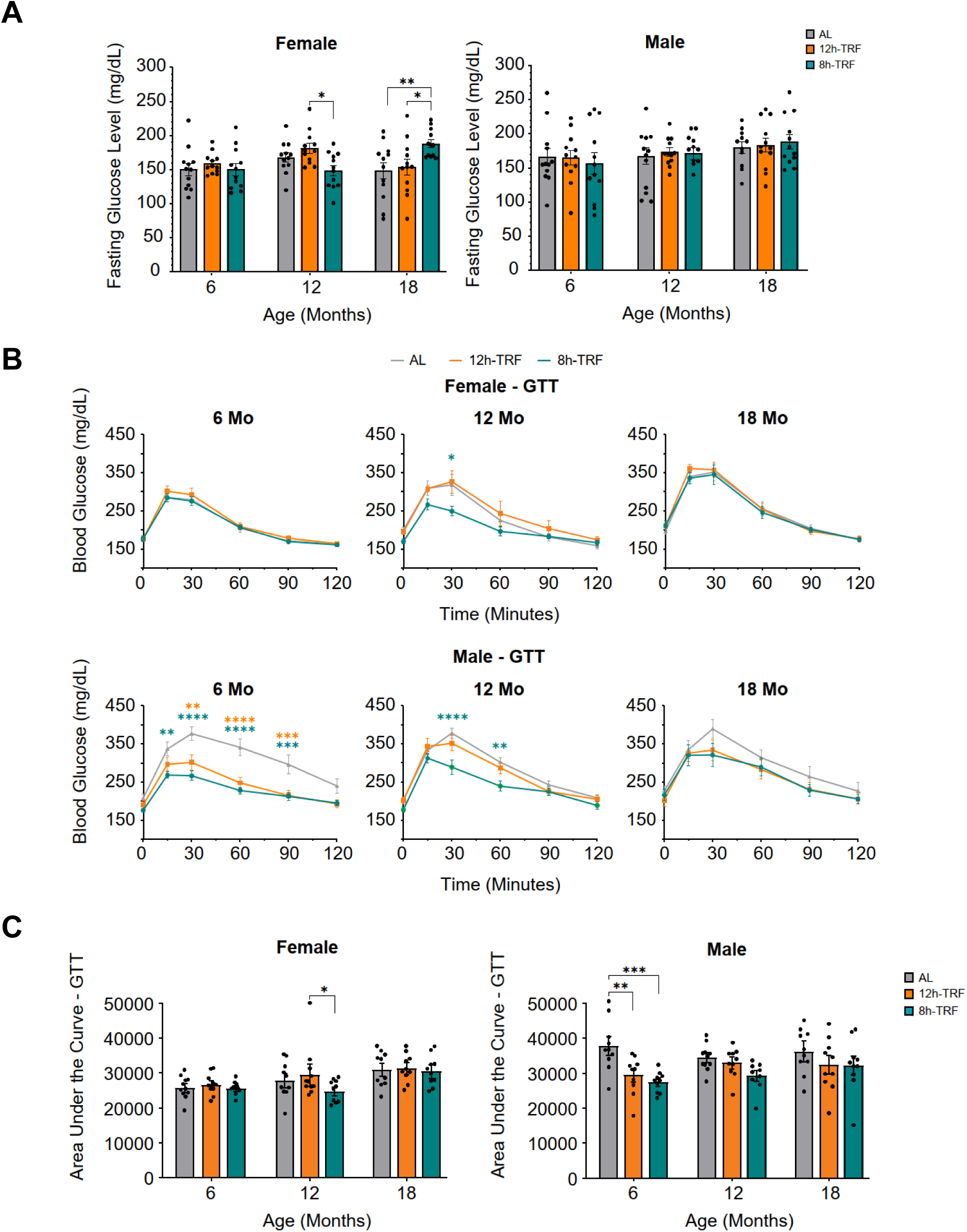
Fasting glucose levels and glucose tolerance tests in follow-up mouse cohort: In females and males: (**A**) Bar graphs of 12h fasting glucose levels (mg/dL) collected at zeitgeber time (ZT) 10, (**B**) glucose tolerance tests (GTT) performed starting at ZT10, and (**C**) area under the curve (AUC) for the GTTs tested at 6, 12, and 18 months of age. Means ± SEM (bars). Two-way ANOVA, Tukey’s post-hoc. *P ≤ 0.05; **P ≤ 0.01; ***P ≤ 0.001; ****P ≤0.0001. N=9-12 per group and sex.

**Figure S10:**
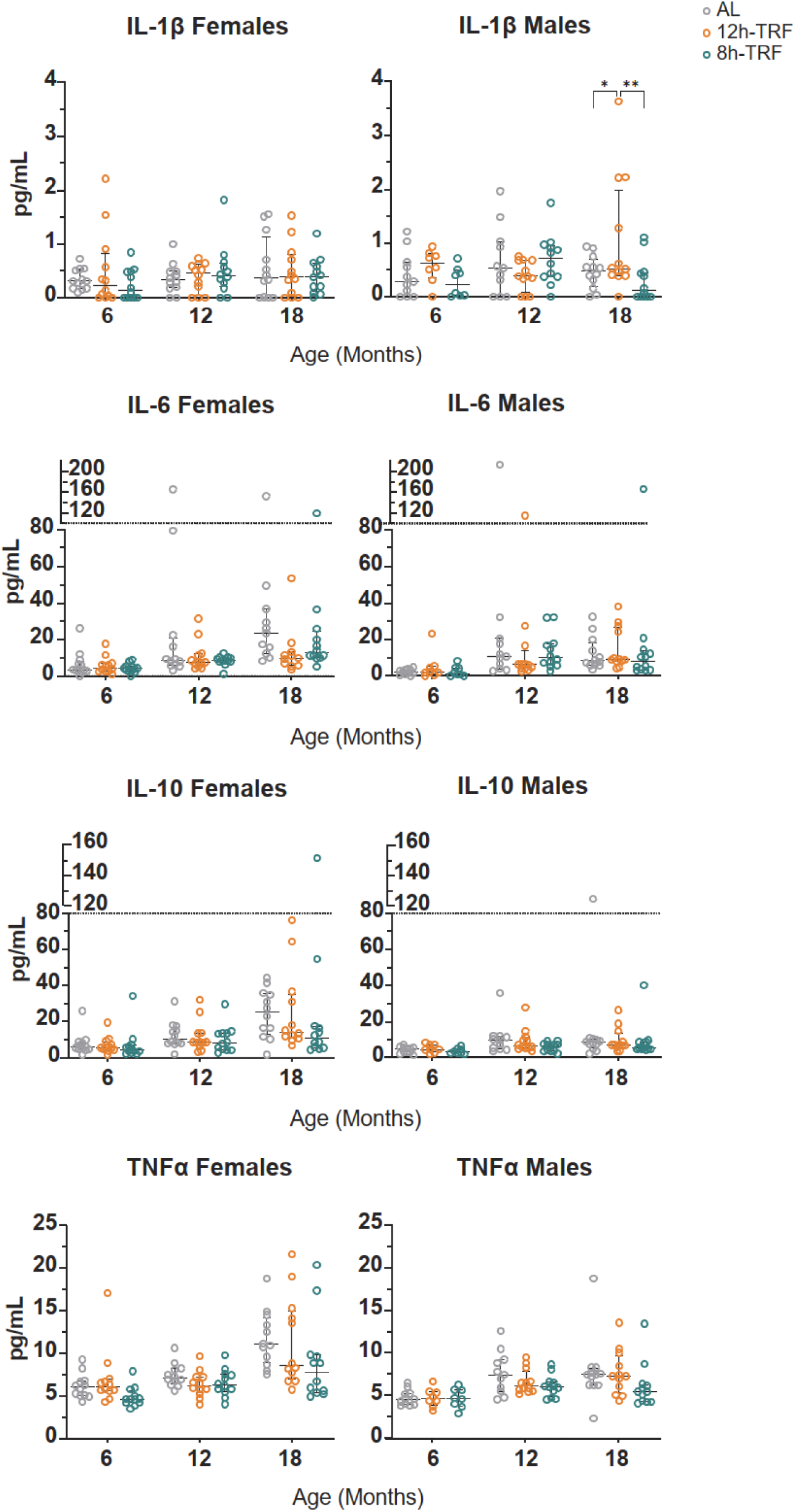
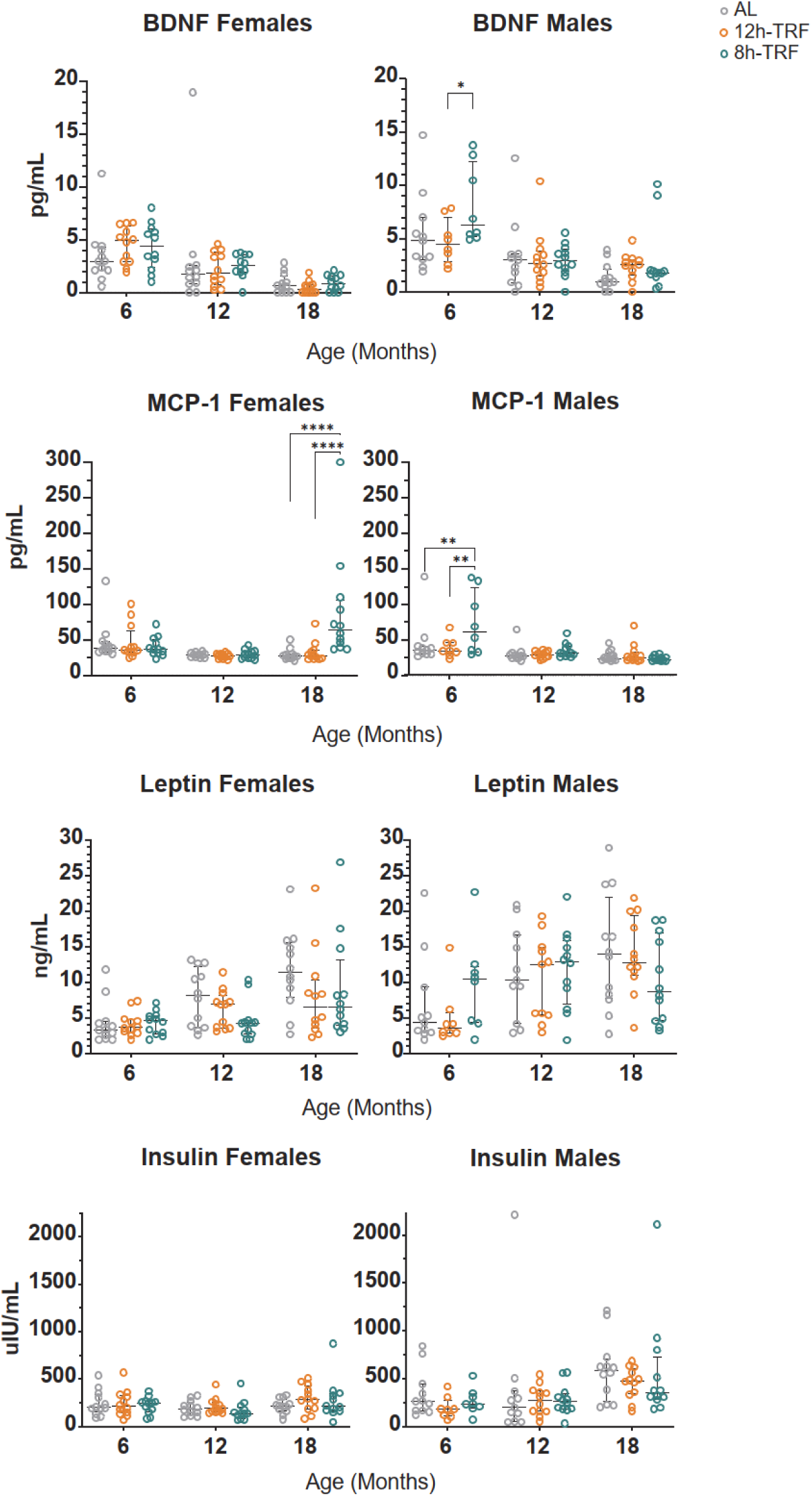
Circulating levels of adipokines from blood plasma in follow-up mouse cohort. In females and males, dot plots of circulating levels of interleukin (IL)-1β, IL-6, IL-10, tumor necrosis factor α (TNFα), brain-derived neurotrophic factor (BDNF), monocyte chemoattractant protein (MCP)-1, leptin, and insulin. Plasma was collected at ZT10 from 12h fasted blood at 6, 12, and 18 months of age. Median with interquartile range. Two-way ANOVA, Tukey’s post-hoc. *P ≤ 0.05; **P ≤ 0.01; ***P ≤ 0.001; ****P ≤0.0001. N=8-12 per group and sex.

**Figure S11:**
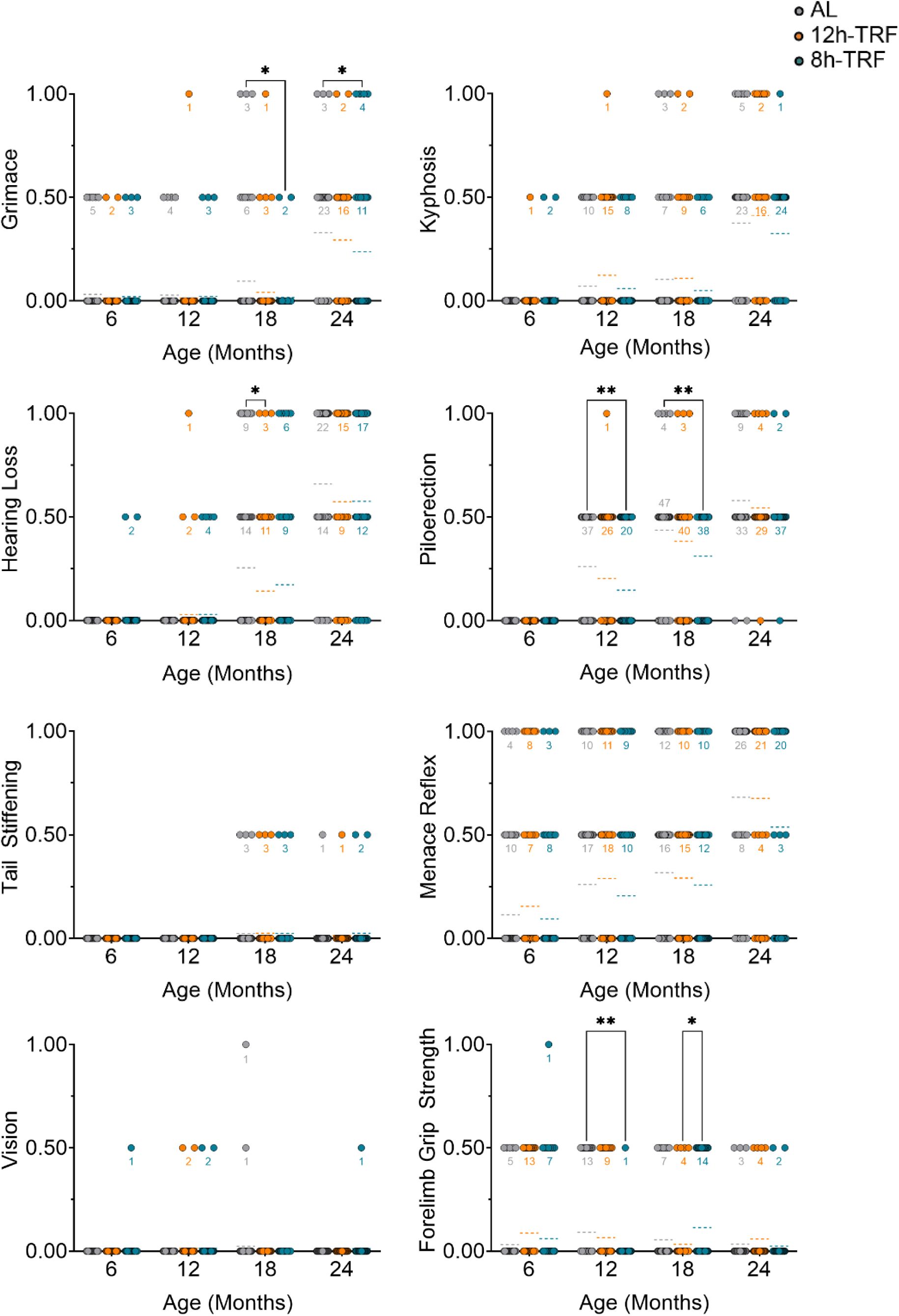
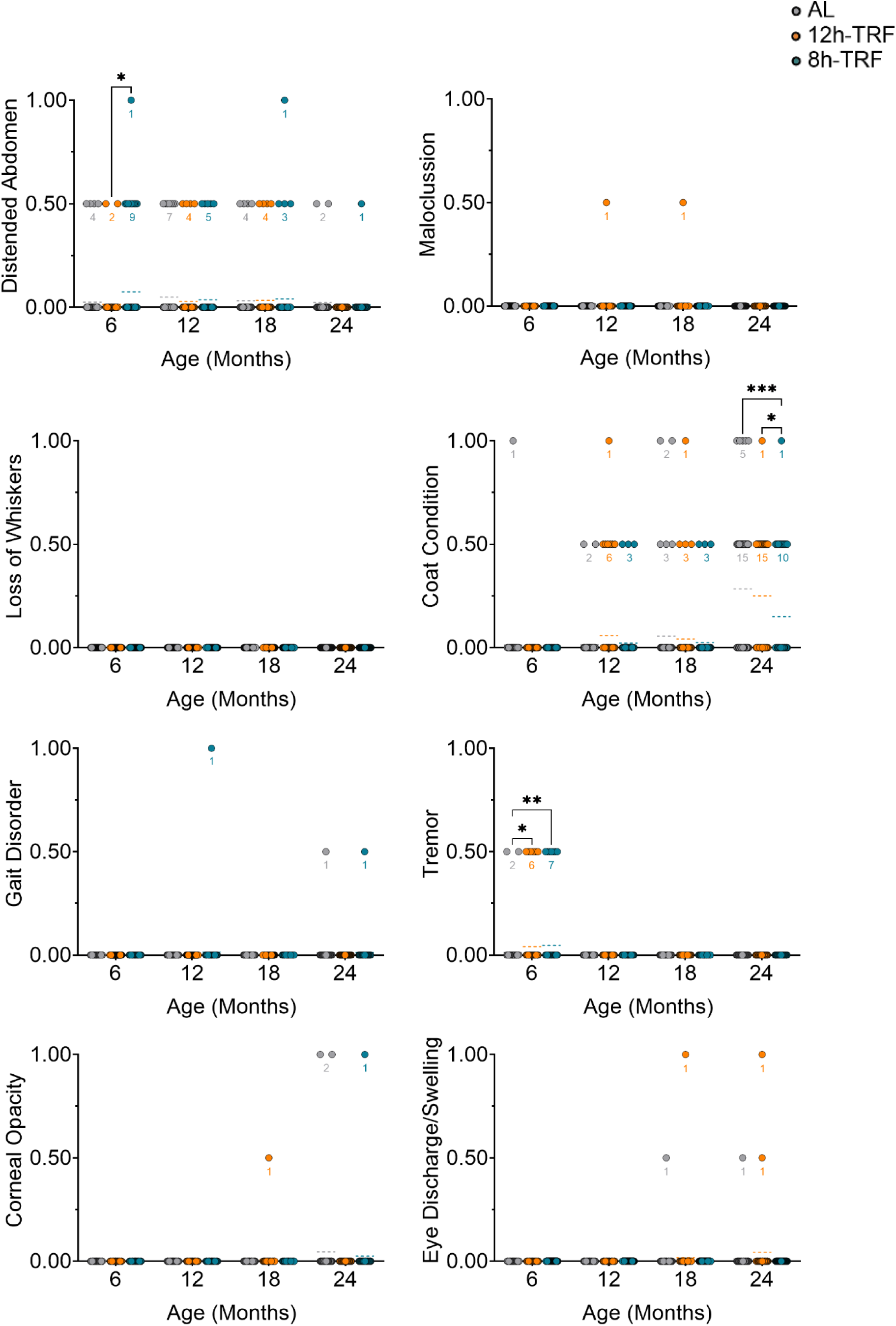
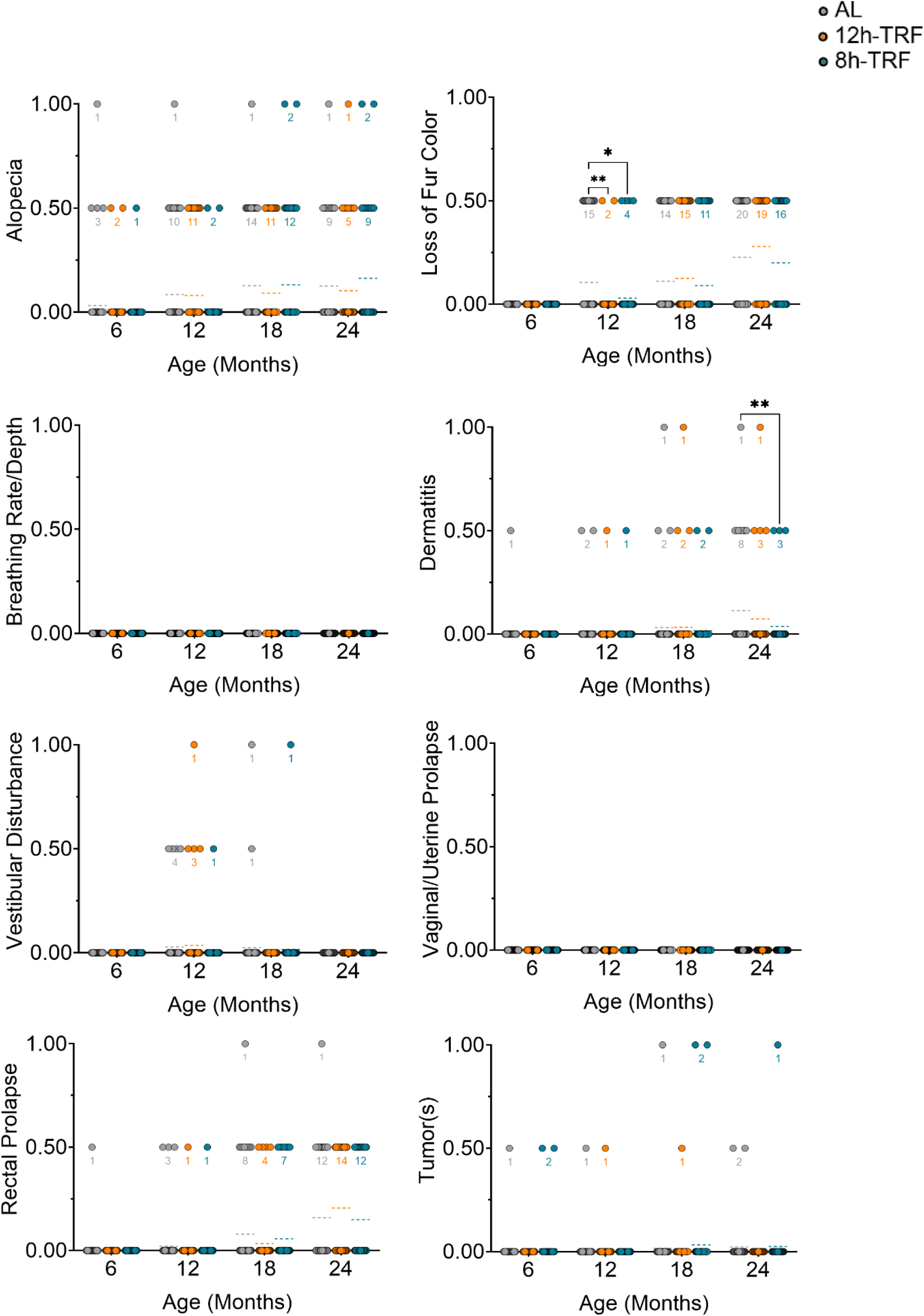
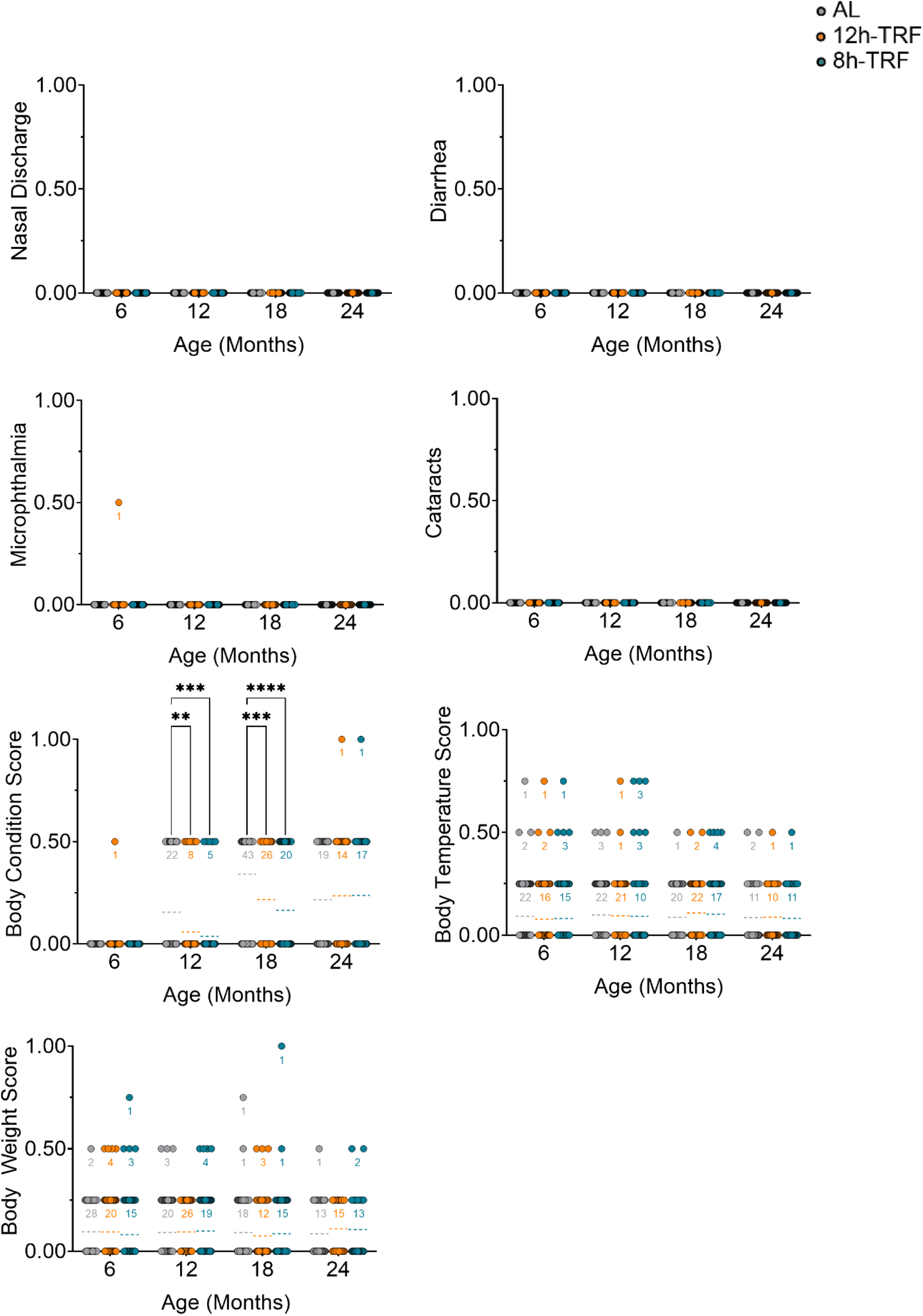
Individual frailty parameter scores in females. Dot plots of frailty scores every 6 months of age in females for each of the 31 parameters modified from (*55*). Most parameters scored as 0 for no, 0.5 for mild, and 1 for severe deficit. Body temperature and body weight scores were scored 0, .25, .5, .75, or 1 with higher values representing more severe deviation of the parameter from the group mean. N values shown in graph for any mice with scores >0. Mean shown as dashed line. Two-way ANOVA, Tukey’s post-hoc. *P ≤ 0.05; **P ≤ 0.01; ***P ≤ 0.001; ****P ≤0.0001. Total N: AL, N=44-79. 12h-TRF, N=34-74. 8h-TRF, N=40-74.

**Figure S12:**
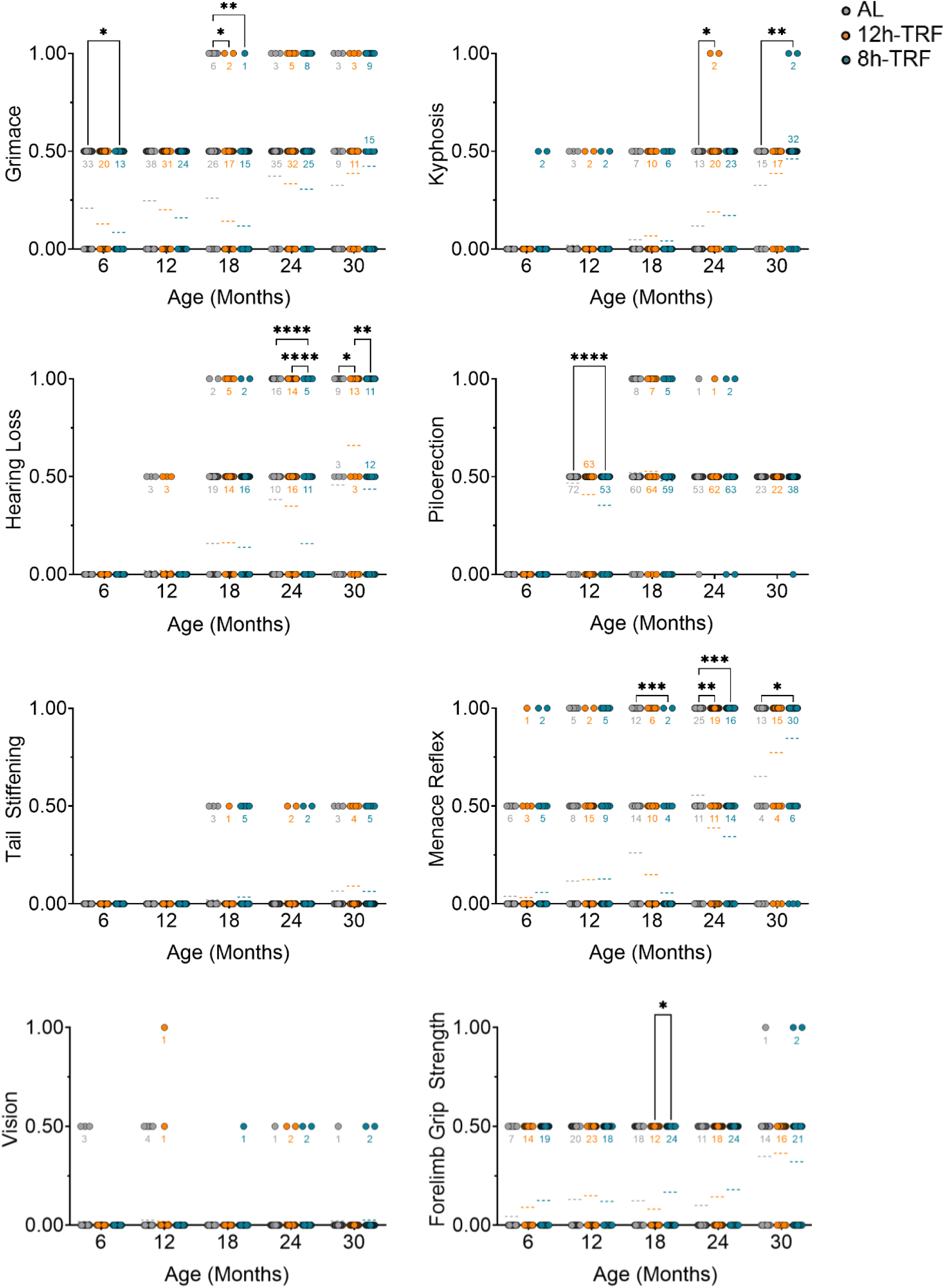
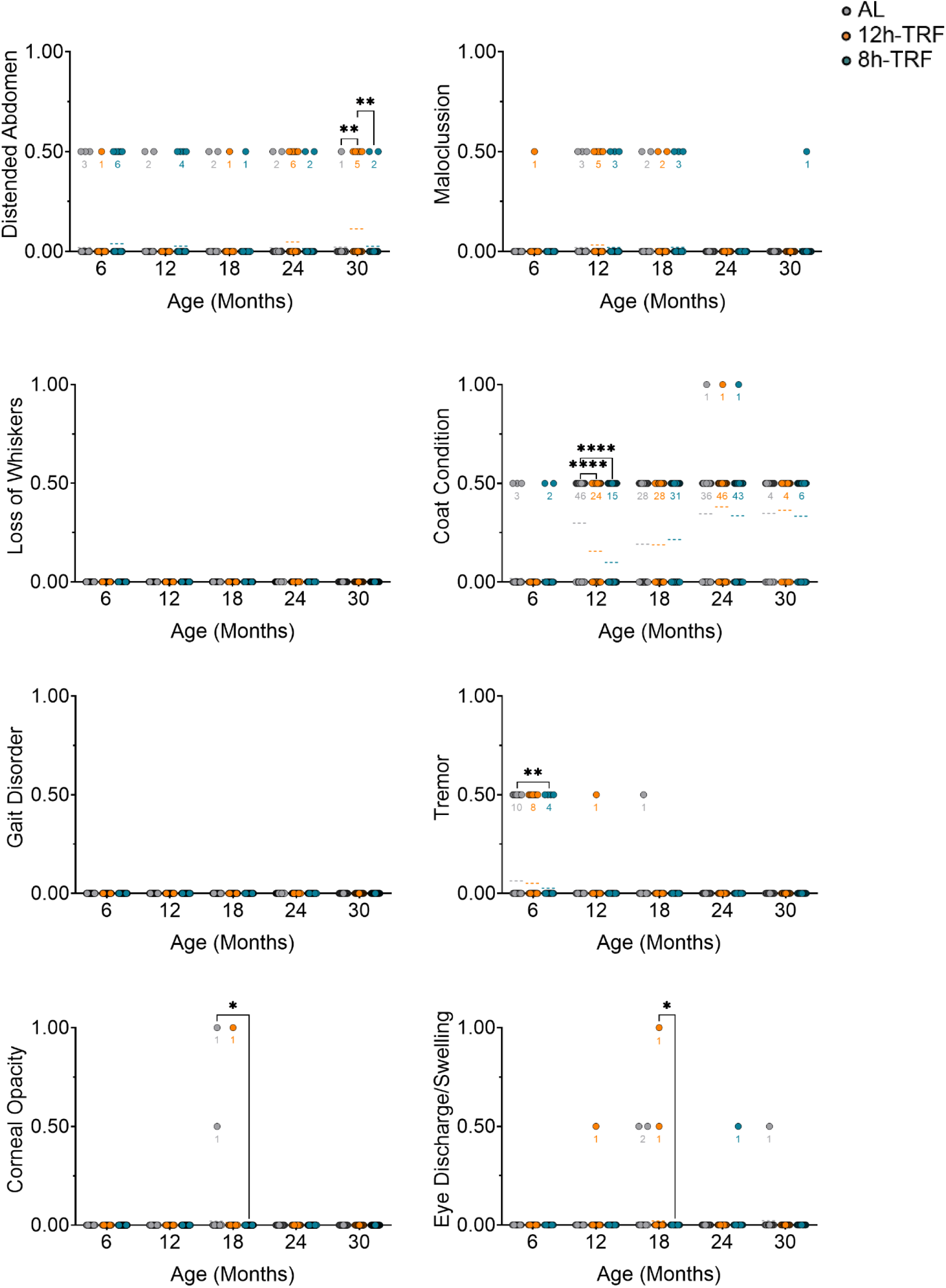
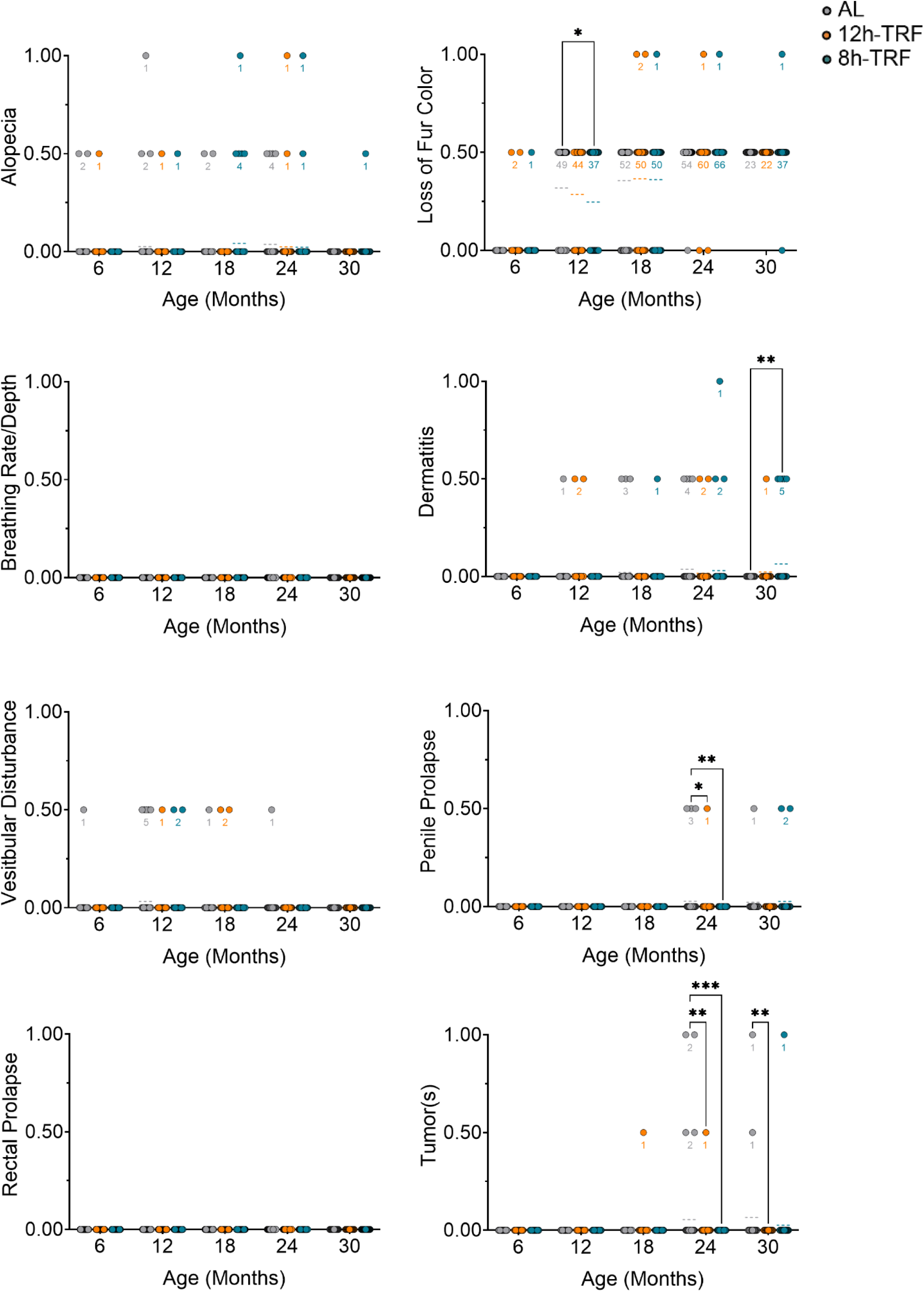
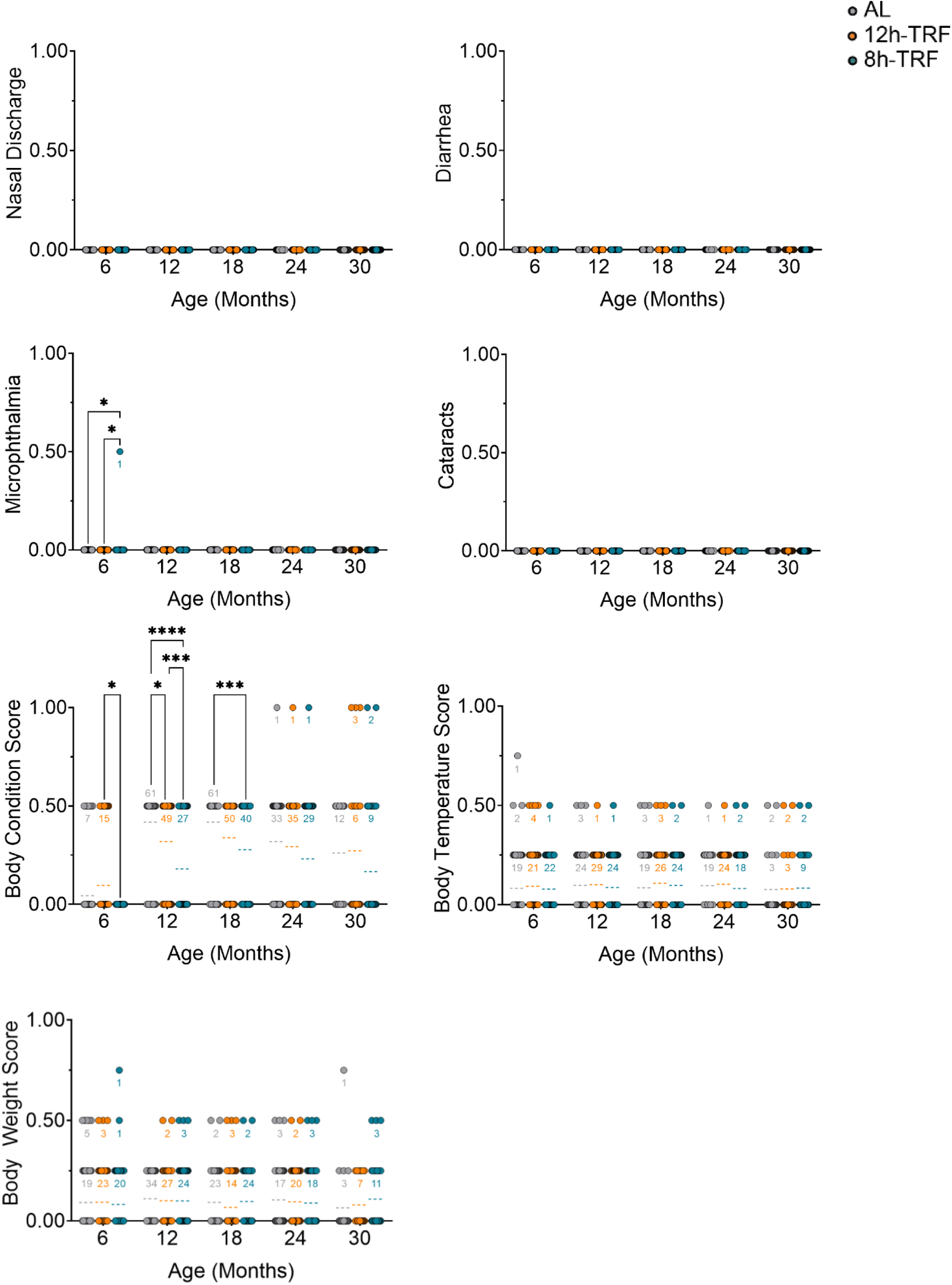
Individual frailty parameter scores in males. Dot plots of frailty scores every 6 months of age in males for each of the 31 parameters modified from (*55*). Most parameters scored as 0 for no, 0.5 for mild, and 1 for severe deficit. Body temperature and body weight scores were scored 0, .25, .5, .75, or 1 with higher values representing more severe deviation of the parameter from the group mean. N values shown in graph for any mice with scores >0. Mean shown as dashed line. Two-way ANOVA, Tukey’s post-hoc. *P ≤ 0.05; **P ≤ 0.01; ***P ≤ 0.001; ****P ≤0.0001. Total N: AL, N=23-79. 12h-TRF, N=22-78. 8h-TRF, N=39-76.

**Figure S13:**
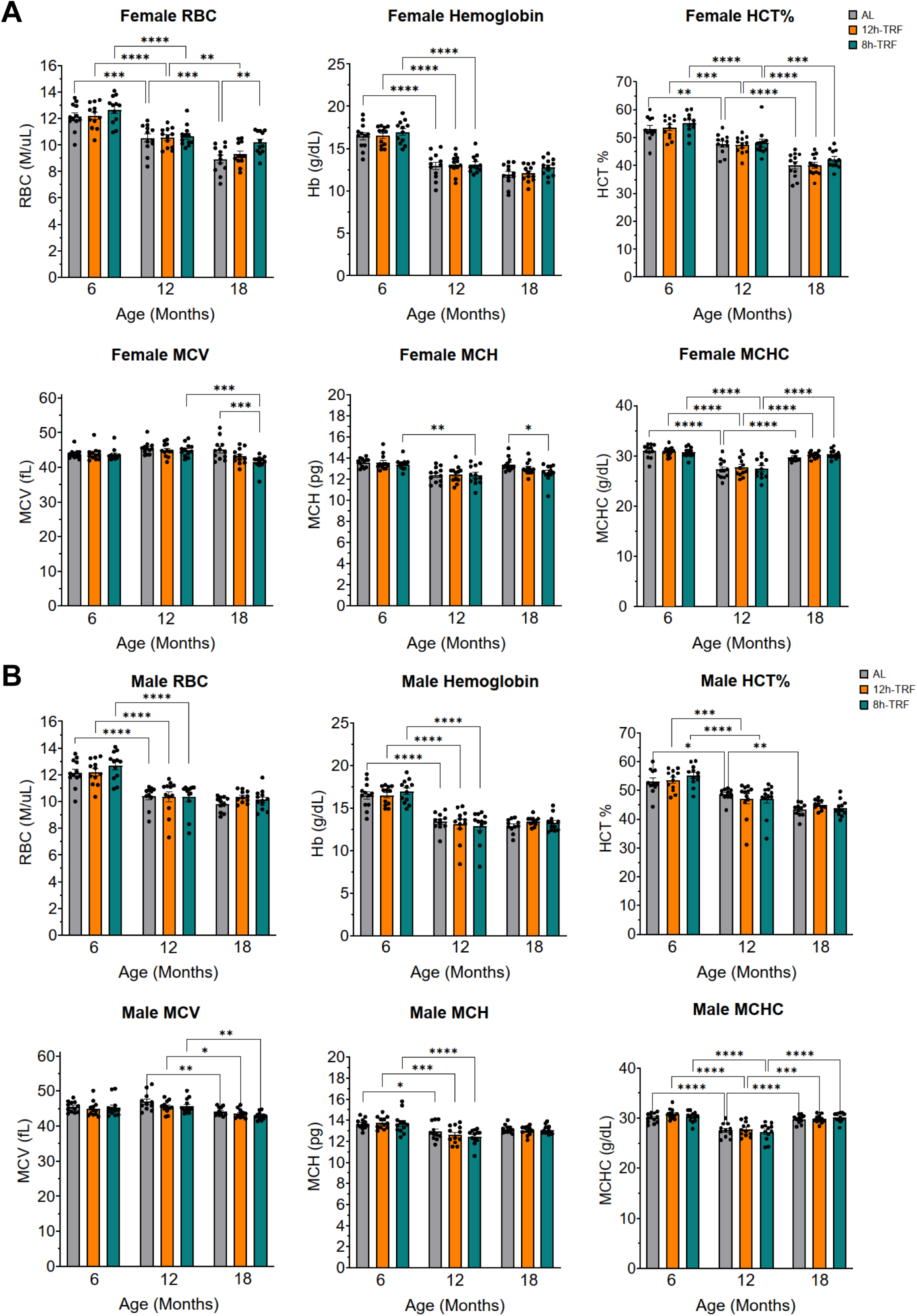
Red blood cell measures with age in follow-up mouse cohort. **(A)** In females and (**B**) males, bar graphs of red blood cell (RBC) counts (M/µL), hemoglobin levels (g/dL), hematocrit (HCT, percentage of RBCs in total blood volume) (%), mean corpuscular volume (MCV, average size of RBCs) (fL), mean corpuscular hemoglobin (MCH, average amount of hemoglobin per RBC) (pg), and mean corpuscular hemoglobin concentration (MCHC, average concentration of hemoglobin within the volume of the RBC) (g/dL). Measured at ZT10 from 12h fasted blood at 6, 12, and 18 months of age. Mean ± SEM (bars). Two-way ANOVA, Tukey’s post-hoc. *P ≤ 0.05; **P ≤ 0.01; ***P ≤ 0.001; ****P ≤0.0001. N=11-12 per group and sex.

**Figure S14:**
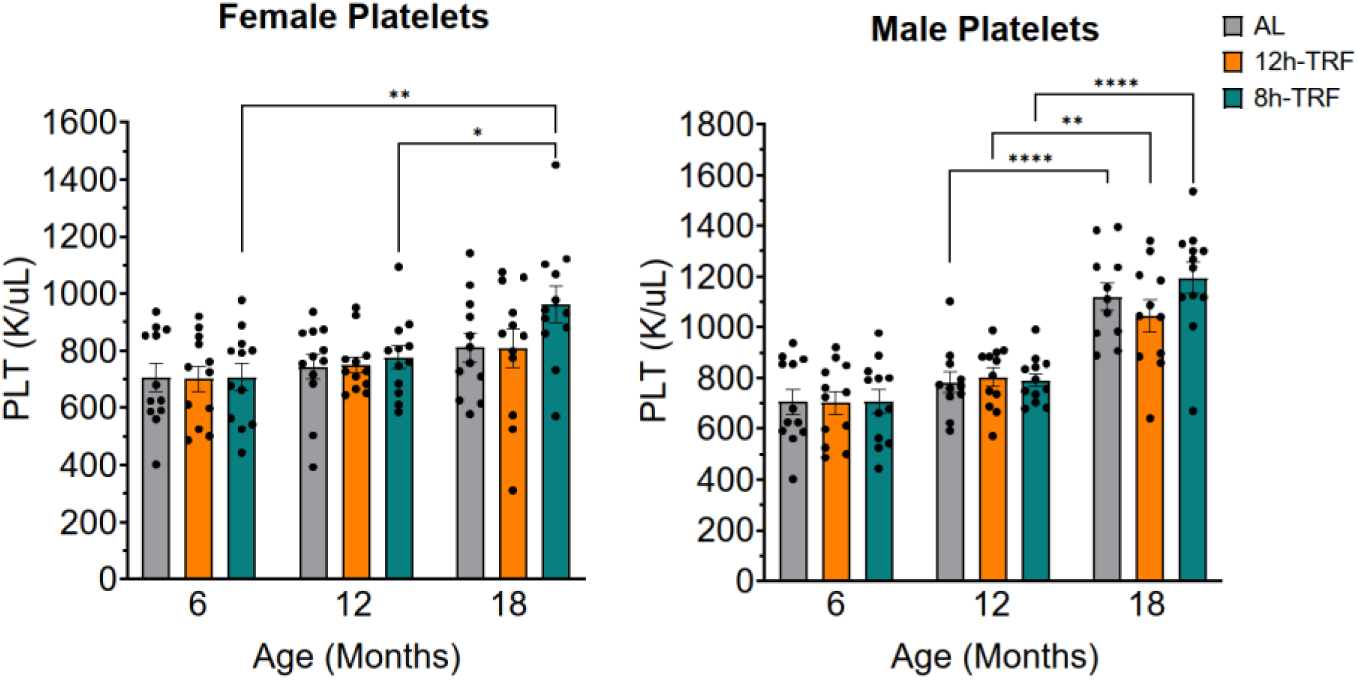
Platelet counts with age in follow-up mouse cohort. In females and males, platelet counts (K/µL). Measured at ZT10 from 12h fasted blood at 6, 12, and 18 months of age. Mean ± SEM (bars). Two-way ANOVA, Tukey’s post-hoc. *P ≤ 0.05; **P ≤ 0.01; ***P ≤ 0.001; ****P ≤0.0001. N=11-12 per group and sex.

**Figure S15:**
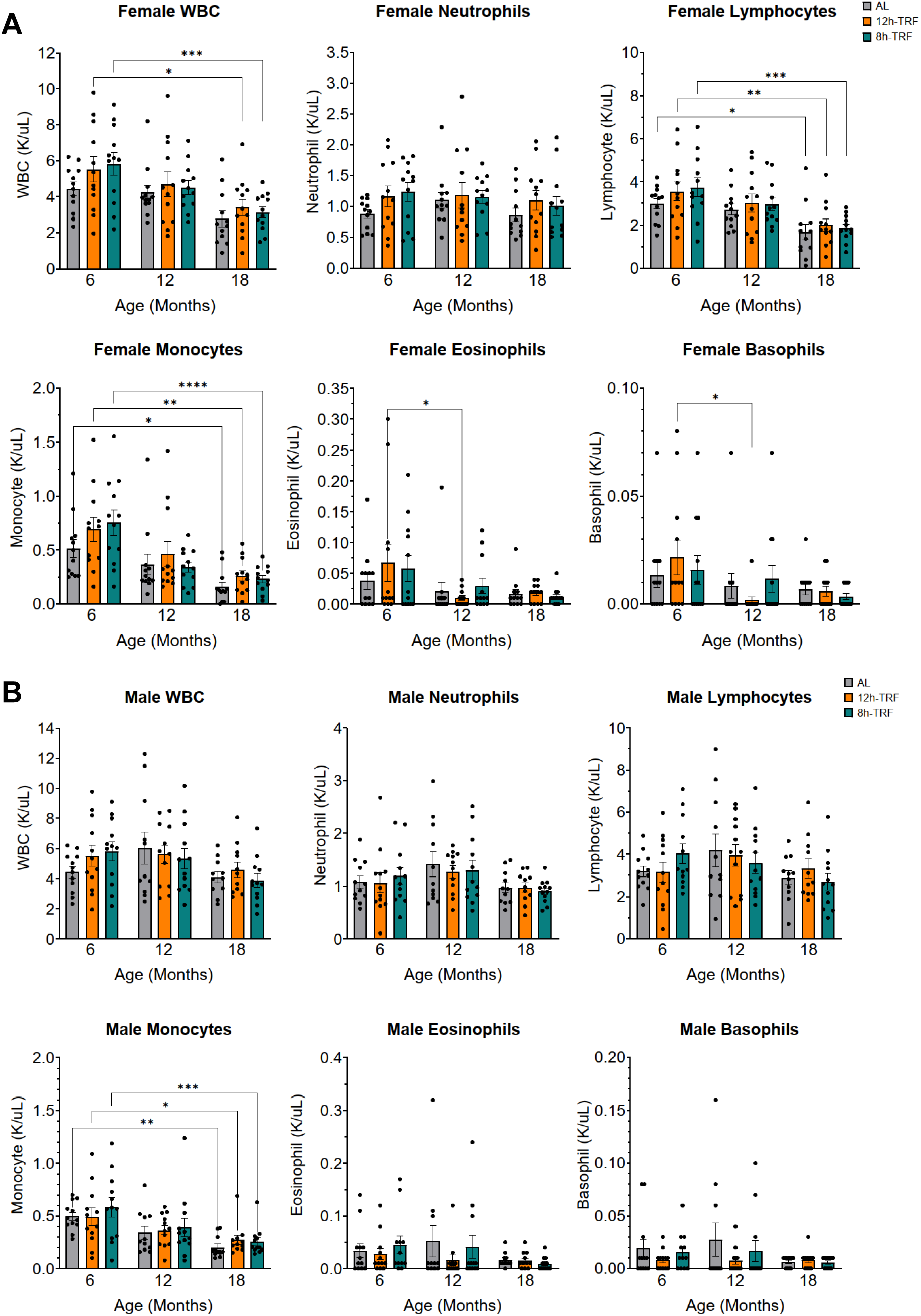
White blood cells measures with age in follow up mouse cohort. (**A**) In females and (**B**) males, bar graphs of total white blood cell (WBC) counts (K/µL) and counts for each WBC sub-type including neutrophils, lymphocytes, monocytes, eosinophils, and basophils. Measured at ZT10 from 12h fasted blood at 6, 12, and 18 months of age. Mean ± SEM (bars). Two-way ANOVA, Tukey’s post-hoc. *P ≤ 0.05; **P ≤ 0.01; ***P ≤ 0.001; ****P ≤0.0001. N=11-12 per group and sex.

**Figure S16:**
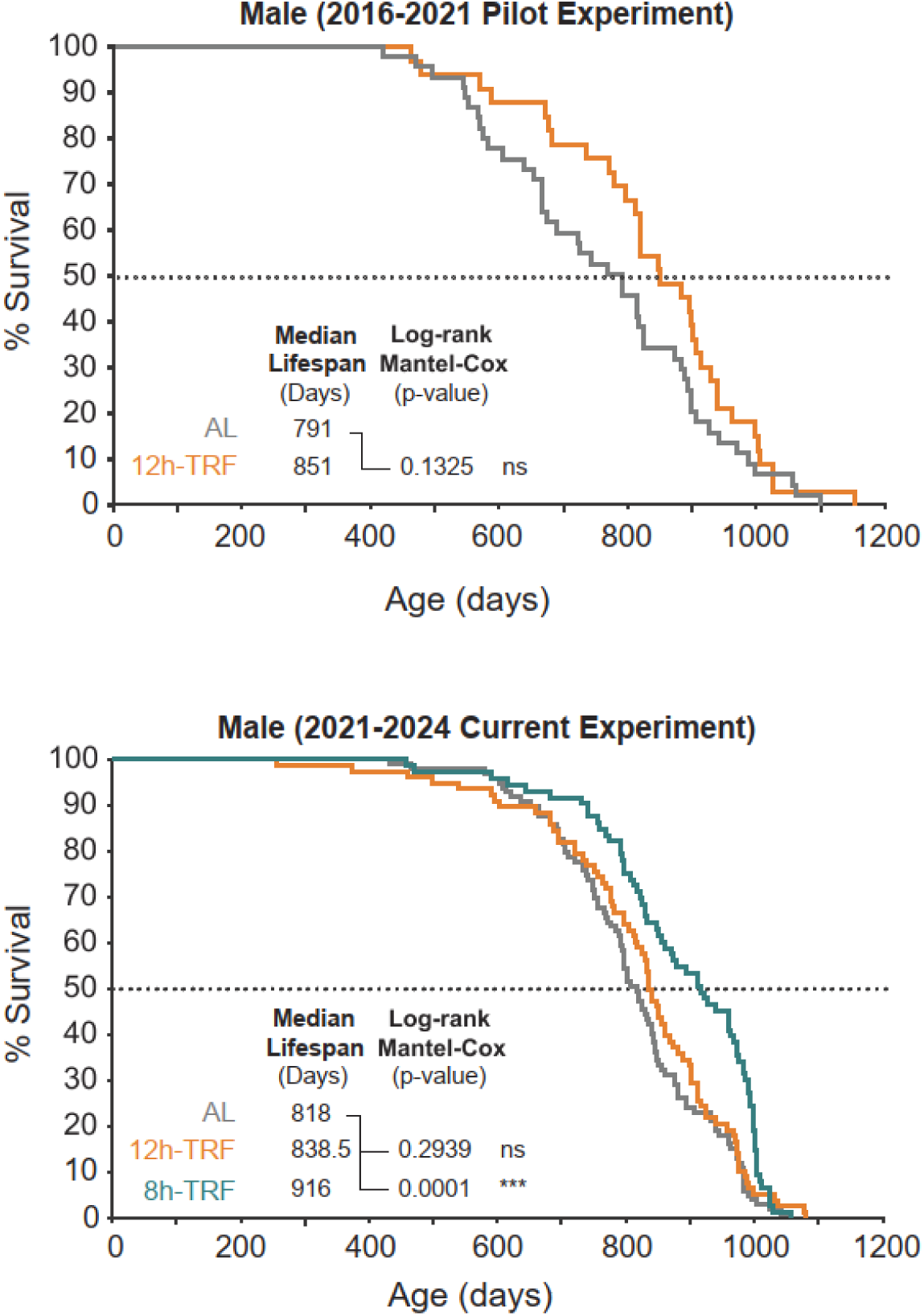
Repeat experiments show 12h-TRF does not extend lifespan in males: Kaplan-Meier survival curves and day median lifespan reached in a pilot experiment vs our current experiment. Log-Rank Mantel-Cox Test for significant difference in overall survival TRF vs AL. Pilot experiment: AL, N=43. 12h-TRF, N=33. Current experiment: AL, N=99. 12h-TRF, N=78. 8h-TRF, N=73.

**Figure S17:**
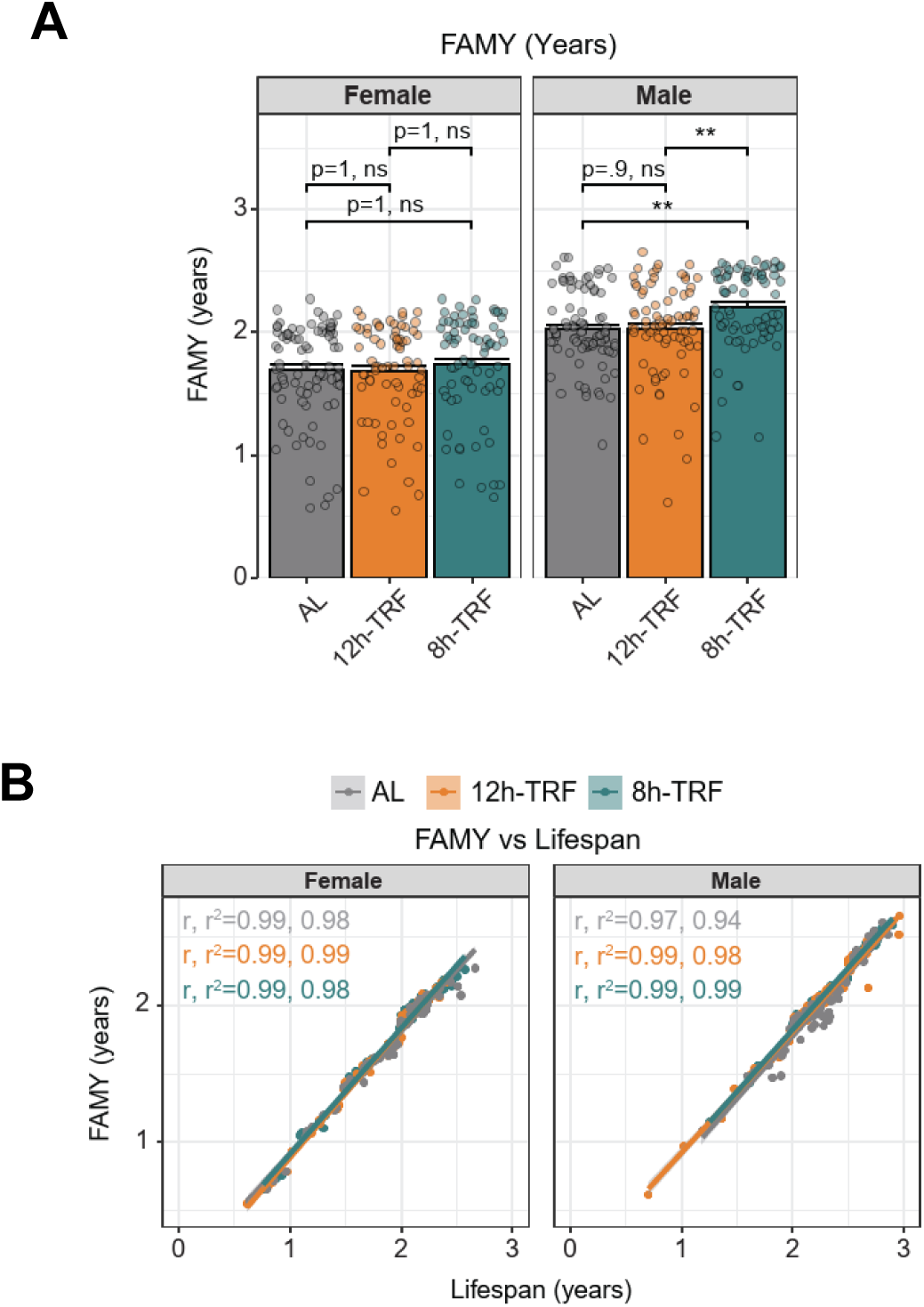
Healthspan as measured by Frailty-Adjusted Mouse Years (FAMY). In females and males: (**A**) FAMY (years) is another method to measure healthspan which utilizes frailty scores as described in (*59*). Higher FAMY = more healthy years. Mean ± SEM (bars). Type III ANOVA, Holm’s post-hoc. *P ≤ 0.05; **P ≤ 0.01; ***P ≤ 0.001; ****P ≤0.0001. (**B**) Correlation plots of FAMY vs lifespan in years. Correlation coefficients (r) and coefficients of determination shown (r^2^).

## Notes

### Competing Interest Statement

The authors have declared no competing interest.

### Summary of Updates

Supplemental files and main text updated with additional follow-up metabolic and inflammatory marker experiments and analyses. Healthspan analysis updated to show TRF extends healthspan in both sexes, but is more prolonged in females. Lifespan analysis was further refined and clarified to show that 8 h-TRF extends lifespan in males only, with no statistically significant impacts in females.

## References and notes

1. D. Costa, Health and the Economy in the United States, from 1750 to the Present. J Econ Lit 53, 503–570 (2015).

2. J. W. Vaupel, Biodemography of human ageing. Nature 464, 536–542 (2010).

3. C. L. Green, D. W. Lamming, L. Fontana, Molecular mechanisms of dietary restriction promoting health and longevity. Nat Rev Mol Cell Biol 23, 56–73 (2022).

4. A. McKay et al., An automated feeding system for the African killifish reveals the impact of diet on lifespan and allows scalable assessment of associative learning. Elife 11, (2022).

5. A. Di Francesco et al., Dietary restriction impacts health and lifespan of genetically diverse mice. Nature 634, 684–692 (2024).

6. S. J. Mitchell et al., Daily Fasting Improves Health and Survival in Male Mice Independent of Diet Composition and Calories. Cell Metab 29, 221–228 e223 (2019).

7. H. H. Pak et al., Fasting drives the metabolic, molecular and geroprotective effects of a calorie-restricted diet in mice. Nat Metab 3, 1327–1341 (2021).

8. E. Duregon et al., Prolonged fasting times reap greater geroprotective effects when combined with caloric restriction in adult female mice. Cell Metab 35, 1179–1194 e1175 (2023).

9. V. A. Acosta-Rodriguez, M. H. M. de Groot, F. Rijo-Ferreira, C. B. Green, J. S. Takahashi, Mice under Caloric Restriction Self-Impose a Temporal Restriction of Food Intake as Revealed by an Automated Feeder System. Cell Metab 26, 267–277 e262 (2017).

10. V. Acosta-Rodriguez et al., Circadian alignment of early onset caloric restriction promotes longevity in male C57BL/6J mice. Science 376, 1192–1202 (2022).

11. H. Reinke, G. Asher, Crosstalk between metabolism and circadian clocks. Nat Rev Mol Cell Biol 20, 227–241 (2019).

12. I. Laothamatas, E. S. Rasmussen, C. B. Green, J. S. Takahashi, Metabolic and chemical architecture of the mammalian circadian clock. Cell Chem Biol 30, 1033–1052 (2023).

13. J. S. Takahashi, Transcriptional architecture of the mammalian circadian clock. Nat Rev Genet 18, 164–179 (2017).

14. G. Manella et al., The liver-clock coordinates rhythmicity of peripheral tissues in response to feeding. Nat Metab 3, 829–842 (2021).

15. B. J. Greenwell et al., Rhythmic Food Intake Drives Rhythmic Gene Expression More Potently than the Hepatic Circadian Clock in Mice. Cell Rep 27, 649–657 e645 (2019).

16. C. Vollmers et al., Time of feeding and the intrinsic circadian clock drive rhythms in hepatic gene expression. Proc Natl Acad Sci U S A 106, 21453–21458 (2009).

17. K. A. Stokkan, S. Yamazaki, H. Tei, Y. Sakaki, M. Menaker, Entrainment of the circadian clock in the liver by feeding. Science 291, 490–493 (2001).

18. L. Palluth, J. S. Takahashi, C. B. Green, Keeping up with the nicotinamides: NADP(H), the forgotten circadian cofactor that keeps metabolic time. Life Metabolism, loaf034 (2025).

19. S. Hood, S. Amir, The aging clock: circadian rhythms and later life. J Clin Invest 127, 437–446 (2017).

20. V. A. Acosta-Rodriguez, F. Rijo-Ferreira, C. B. Green, J. S. Takahashi, Importance of circadian timing for aging and longevity. Nat Commun 12, 2862 (2021).

21. A. Chaix, E. N. C. Manoogian, G. C. Melkani, S. Panda, Time-Restricted Eating to Prevent and Manage Chronic Metabolic Diseases. Annu Rev Nutr 39, 291–315 (2019).

22. W. E. Kraus et al., 2 years of calorie restriction and cardiometabolic risk (CALERIE): exploratory outcomes of a multicentre, phase 2, randomised controlled trial. Lancet Diabetes Endocrinol 7, 673–683 (2019).

23. V. D. Longo, S. Panda, Fasting, Circadian Rhythms, and Time-Restricted Feeding in Healthy Lifespan. Cell Metab 23, 1048–1059 (2016).

24. M. E. Parrotta et al., Time Restricted Eating: A Valuable Alternative to Calorie Restriction for Addressing Obesity? Curr Obes Rep 14, 17 (2025).

25. D. M. Arble, J. Bass, A. D. Laposky, M. H. Vitaterna, F. W. Turek, Circadian timing of food intake contributes to weight gain. Obesity (Silver Spring) 17, 2100–2102 (2009).

26. E. N. C. Manoogian, L. S. Chow, P. R. Taub, B. Laferrere, S. Panda, Time-restricted Eating for the Prevention and Management of Metabolic Diseases. Endocr Rev 43, 405–436 (2022).

27. P. Balasubramanian et al., Time-restricted feeding (TRF) for prevention of age-related vascular cognitive impairment and dementia. Ageing Res Rev 64, 101189 (2020).

28. S. Deota et al., Diurnal transcriptome landscape of a multi-tissue response to time-restricted feeding in mammals. Cell Metab 35, 150–165 e154 (2023).

29. M. Hatori et al., Time-restricted feeding without reducing caloric intake prevents metabolic diseases in mice fed a high-fat diet. Cell Metab 15, 848–860 (2012).

30. H. Sherman et al., Timed high-fat diet resets circadian metabolism and prevents obesity. FASEB J 26, 3493–3502 (2012).

31. P. Regmi et al., Early or delayed time-restricted feeding prevents metabolic impact of obesity in mice. J Endocrinol 248, 75–86 (2021).

32. C. Hepler et al., Time-restricted feeding mitigates obesity through adipocyte thermogenesis. Science 378, 276–284 (2022).

33. E. N. C. Manoogian et al., Feasibility of time-restricted eating and impacts on cardiometabolic health in 24-h shift workers: The Healthy Heroes randomized control trial. Cell Metab 34, 1442–1456 e1447 (2022).

34. K. L. Haganes et al., Time-restricted eating and exercise training improve HbA1c and body composition in women with overweight/obesity: A randomized controlled trial. Cell Metab 34, 1457–1471 e1454 (2022).

35. T. Che et al., Time-restricted feeding improves blood glucose and insulin sensitivity in overweight patients with type 2 diabetes: a randomised controlled trial. Nutr Metab (Lond) 18, 88 (2021).

36. M. Dote-Montero et al., Effects of early, late and self-selected time-restricted eating on visceral adipose tissue and cardiometabolic health in participants with overweight or obesity: a randomized controlled trial. Nat Med 31, 524–533 (2025).

37. V. Pavlou et al., Effect of Time-Restricted Eating on Weight Loss in Adults With Type 2 Diabetes: A Randomized Clinical Trial. JAMA Netw Open 6, e2339337 (2023).

38. Z. Xie et al., Randomized controlled trial for time-restricted eating in healthy volunteers without obesity. Nat Commun 13, 1003 (2022).

39. A. Chaix, A. Zarrinpar, P. Miu, S. Panda, Time-restricted feeding is a preventative and therapeutic intervention against diverse nutritional challenges. Cell Metab 20, 991–1005 (2014).

40. D. S. Whittaker et al., Circadian modulation by time-restricted feeding rescues brain pathology and improves memory in mouse models of Alzheimer’s disease. Cell Metab 35, 1704–1721 e1706 (2023).

41. A. Chaix, S. Deota, R. Bhardwaj, T. Lin, S. Panda, Sex– and age-dependent outcomes of 9-hour time-restricted feeding of a Western high-fat high-sucrose diet in C57BL/6J mice. Cell Rep 36, 109543 (2021).

42. M. Ulgherait et al., Circadian autophagy drives iTRF-mediated longevity. Nature 598, 353–358 (2021).

43. K. Xie et al., Every-other-day feeding extends lifespan but fails to delay many symptoms of aging in mice. Nat Commun 8, 155 (2017).

44. M. E. Starr, H. Saito, Age-related increase in food spilling by laboratory mice may lead to significant overestimation of actual food consumption: implications for studies on dietary restriction, metabolism, and dose calculations. J Gerontol A Biol Sci Med Sci 67, 1043–1048 (2012).

45. Y. D. Rathod, M. Di Fulvio, The feeding microstructure of male and female mice. PLoS One 16, e0246569 (2021).

46. K. Gabel et al., Effects of 8-hour time restricted feeding on body weight and metabolic disease risk factors in obese adults: A pilot study. Nutr Healthy Aging 4, 345–353 (2018).

47. V. A. Acosta-Rodriguez et al., Misaligned feeding uncouples daily rhythms within brown adipose tissue and between peripheral clocks. Cell Rep 43, 114523 (2024).

48. G. Manzanares, G. Brito-da-Silva, P. G. Gandra, Voluntary wheel running: patterns and physiological effects in mice. Braz J Med Biol Res 52, e7830 (2018).

49. V. S. Valentinuzzi, K. Scarbrough, J. S. Takahashi, F. W. Turek, Effects of aging on the circadian rhythm of wheel-running activity in C57BL/6 mice. Am J Physiol 273, R1957–1964 (1997).

50. R. Garcia-Valles et al., Life-long spontaneous exercise does not prolong lifespan but improves health span in mice. Longev Healthspan 2, 14 (2013).

51. O. Froy, Circadian aspects of energy metabolism and aging. Ageing Res Rev 12, 931–940 (2013).

52. C. Lopez-Otin, L. Galluzzi, J. M. P. Freije, F. Madeo, G. Kroemer, Metabolic Control of Longevity. Cell 166, 802–821 (2016).

53. D. Gatfield, U. Schibler, Circadian glucose homeostasis requires compensatory interference between brain and liver clocks. Proc Natl Acad Sci U S A 105, 14753–14754 (2008).

54. D. L. Palliyaguru et al., Fasting blood glucose as a predictor of mortality: Lost in translation. Cell Metab 33, 2189–2200 e2183 (2021).

55. J. C. Whitehead et al., A clinical frailty index in aging mice: comparisons with frailty index data in humans. J Gerontol A Biol Sci Med Sci 69, 621–632 (2014).

56. J. Martinez-Romero et al., A hematology-based clock derived from the Study of Longitudinal Aging in Mice to estimate biological age. Nat Aging 4, 1882–1896 (2024).

57. M. Magnani et al., Effect of age on some properties of mice erythrocytes. Mechanisms of Ageing and Development 42, 37–47 (1988).

58. D. L. Palliyaguru, J. M. Moats, C. Di Germanio, M. Bernier, R. de Cabo, Frailty index as a biomarker of lifespan and healthspan: Focus on pharmacological interventions. Mech Ageing Dev 180, 42–48 (2019).

59. D. W. Lamming, Quantification of healthspan in aging mice: introducing FAMY and GRAIL. Geroscience 46, 4203–4215 (2024).

60. R. Huang et al., Multi-omics profiling reveals rhythmic liver function shaped by meal timing. Nat Commun 14, 6086 (2023).

61. J. S. Pendergast, K. L. Branecky, R. Huang, K. D. Niswender, S. Yamazaki, Wheel-running activity modulates circadian organization and the daily rhythm of eating behavior. Front Psychol 5, 177 (2014).

62. A. M. Schroeder et al., Voluntary scheduled exercise alters diurnal rhythms of behaviour, physiology and gene expression in wild-type and vasoactive intestinal peptide-deficient mice. J Physiol 590, 6213–6226 (2012).

63. I. Janssen, S. B. Heymsfield, R. Ross, Low relative skeletal muscle mass (sarcopenia) in older persons is associated with functional impairment and physical disability. J Am Geriatr Soc 50, 889–896 (2002).

64. E. Marzetti, C. Leeuwenburgh, Skeletal muscle apoptosis, sarcopenia and frailty at old age. Exp Gerontol 41, 1234–1238 (2006).

65. C. Livelo, Y. Guo, G. C. Melkani, A Skeletal Muscle-Centric View on Time-Restricted Feeding and Obesity under Various Metabolic Challenges in Humans and Animals. Int J Mol Sci 24, (2022).

66. Y. O. Henderson et al., Late-life intermittent fasting decreases aging-related frailty and increases renal hydrogen sulfide production in a sexually dimorphic manner. Geroscience 43, 1527–1554 (2021).

67. Y. Yamada et al., Caloric Restriction and Healthy Life Span: Frail Phenotype of Nonhuman Primates in the Wisconsin National Primate Research Center Caloric Restriction Study. J Gerontol A Biol Sci Med Sci 73, 273–278 (2018).

68. O. Arum, Z. A. Rasche, D. J. Rickman, A. Bartke, Prevention of neuromusculoskeletal frailty in slow-aging ames dwarf mice: longitudinal investigation of interaction of longevity genes and caloric restriction. PLoS One 8, e72255 (2013).

69. A. E. Kane et al., Impact of Longevity Interventions on a Validated Mouse Clinical Frailty Index. J Gerontol A Biol Sci Med Sci 71, 333–339 (2016).

70. A. E. Kane et al., Animal models of frailty: current applications in clinical research. Clin Interv Aging 11, 1519–1529 (2016).

71. R. Lok, J. Qian, S. L. Chellappa, Sex differences in sleep, circadian rhythms, and metabolism: Implications for precision medicine. Sleep Med Rev 75, 101926 (2024).

72. G. K. Paschos, R. Lordan, G. A. FitzGerald, Intersection of sex and circadian biology. Current Opinion in Physiology 45, 100834 (2025).

73. Z. Bahadoran, P. Mirmiran, K. Kashfi, A. Ghasemi, Effects of time-restricted feeding (TRF)-model of intermittent fasting on adipose organ: a narrative review. Eat Weight Disord 29, 77 (2024).

74. P. Regmi, L. K. Heilbronn, Time-Restricted Eating: Benefits, Mechanisms, and Challenges in Translation. iScience 23, 101161 (2020).

75. P. Chen, B. Li, L. Ou-Yang, Role of estrogen receptors in health and disease. Front Endocrinol (Lausanne) 13, 839005 (2022).

76. V. M. Alvord, E. J. Kantra, J. S. Pendergast, Estrogens and the circadian system. Semin Cell Dev Biol 126, 56–65 (2022).

77. K. M. Hatcher, S. E. Royston, M. M. Mahoney, Modulation of circadian rhythms through estrogen receptor signaling. Eur J Neurosci 51, 217–228 (2020).

78. L. Zhu et al., Estrogens prevent metabolic dysfunctions induced by circadian disruptions in female mice. Endocrinology 156, 2114–2123 (2015).

79. L. Talamanca, C. Gobet, F. Naef, Sex-dimorphic and age-dependent organization of 24-hour gene expression rhythms in humans. Science 379, 478–483 (2023).

80. J. A. Krizo, E. M. Mintz, Sex differences in behavioral circadian rhythms in laboratory rodents. Front Endocrinol (Lausanne) 5, 234 (2014).

81. A. A. Astafev, V. Mezhnina, A. Poe, P. Jiang, R. V. Kondratov, Sexual dimorphism of circadian liver transcriptome. iScience 27, 109483 (2024).

82. S. J. Mitchell et al., Effects of Sex, Strain, and Energy Intake on Hallmarks of Aging in Mice. Cell Metab 23, 1093–1112 (2016).

83. A. E. Kane, D. A. Sinclair, J. R. Mitchell, S. J. Mitchell, Sex differences in the response to dietary restriction in rodents. Curr Opin Physiol 6, 28–34 (2018).

84. J. F. Lemaitre et al., Sex differences in adult lifespan and aging rates of mortality across wild mammals. Proc Natl Acad Sci U S A 117, 8546–8553 (2020).

85. S. N. Austad, K. E. Fischer, Sex Differences in Lifespan. Cell Metab 23, 1022–1033 (2016).

86. D. E. Harrison et al., Acarbose, 17-alpha-estradiol, and nordihydroguaiaretic acid extend mouse lifespan preferentially in males. Aging Cell 13, 273–282 (2014).

87. R. A. Miller et al., Rapamycin, but not resveratrol or simvastatin, extends life span of genetically heterogeneous mice. J Gerontol A Biol Sci Med Sci 66, 191–201 (2011).

88. R. Strong et al., Nordihydroguaiaretic acid and aspirin increase lifespan of genetically heterogeneous male mice. Aging Cell 7, 641–650 (2008).

89. B. N. Gaskill et al., Impact of nesting material on mouse body temperature and physiology. Physiol Behav 110-111, 87–95 (2013).

90. K. Kaikaew, J. Steenbergen, A. P. N. Themmen, J. A. Visser, A. Grefhorst, Sex difference in thermal preference of adult mice does not depend on presence of the gonads. Biol Sex Differ 8, 24 (2017).

91. K. Kaikaew et al., Sex difference in cold perception and shivering onset upon gradual cold exposure. J Therm Biol 77, 137–144 (2018).

92. C. Fernandez-Pena, A. Reimundez, F. Viana, V. M. Arce, R. Senaris, Sex differences in thermoregulation in mammals: Implications for energy homeostasis. Front Endocrinol (Lausanne) 14, 1093376 (2023).

93. Z. Zhao et al., Body temperature is a more important modulator of lifespan than metabolic rate in two small mammals. Nat Metab 4, 320–326 (2022).

94. B. Conti et al., Transgenic mice with a reduced core body temperature have an increased life span. Science 314, 825–828 (2006).

95. A. W. Fischer, B. Cannon, J. Nedergaard, Optimal housing temperatures for mice to mimic the thermal environment of humans: An experimental study. Mol Metab 7, 161–170 (2018).

96. C. L. Karp, Unstressing intemperate models: how cold stress undermines mouse modeling. J Exp Med 209, 1069–1074 (2012).

97. A. L. Ray et al., The role of sex in the innate and adaptive immune environment of metastatic colorectal cancer. Br J Cancer 123, 624–632 (2020).

98. E. G. Vichaya et al., Sex differences in the behavioral and immune responses of mice to tumor growth and cancer therapy. Brain Behav Immun 98, 161–172 (2021).

99. J. B. Rubin, The spectrum of sex differences in cancer. Trends Cancer 8, 303–315 (2022).

100. M. K. E. Ougaard et al., Murine Nephrotoxic Nephritis as a Model of Chronic Kidney Disease. Int J Nephrol 2018, 8424502 (2018).

101. United States Renal Data System. (National Institute of Diabetes and Digestive and Kidney Diseases, National Institutes of Health, Bethesda, MD).

102. G. J. Shim, L. L. Kis, M. Warner, J. A. Gustafsson, Autoimmune glomerulonephritis with spontaneous formation of splenic germinal centers in mice lacking the estrogen receptor alpha gene. Proc Natl Acad Sci U S A 101, 1720–1724 (2004).

103. J. H. Catterson et al., Short-Term, Intermittent Fasting Induces Long-Lasting Gut Health and TOR-Independent Lifespan Extension. Curr Biol 28, 1714–1724 e1714 (2018).

104. Y. Yang, D. Liu, Impacts of time-restricted feeding on middle-aged and old mice with obesity. J Physiol, (2024).

105. P. Domaszewski et al., Effect of a six-week times restricted eating intervention on the body composition in early elderly men with overweight. Sci Rep 12, 9816 (2022).

106. O. Hahn et al., A nutritional memory effect counteracts benefits of dietary restriction in old mice. Nat Metab 1, 1059–1073 (2019).

107. A. C. Hofacker, M. Knop, S. Krauss-Etschmann, T. Roeder, Time-Restricted Feeding Promotes Longevity and Gut Health Without Fitness Trade-Offs. FASEB J 39, e70627 (2025).

108. M. J. Schafer et al., Late-life time-restricted feeding and exercise differentially alter healthspan in obesity. Aging Cell 18, e12966 (2019).

109. M. Das et al., Time-Restricted Feeding Attenuates Metabolic Dysfunction-Associated Steatohepatitis and Hepatocellular Carcinoma in Obese Male Mice. Cancers (Basel) 16, (2024).

110. E. B. Parr, B. L. Devlin, B. E. Radford, J. A. Hawley, A Delayed Morning and Earlier Evening Time-Restricted Feeding Protocol for Improving Glycemic Control and Dietary Adherence in Men with Overweight/Obesity: A Randomized Controlled Trial. Nutrients 12, (2020).

111. K. Gabel, S. Cienfuegos, F. Kalam, M. Ezpeleta, K. A. Varady, Time-Restricted Eating to Improve Cardiovascular Health. Curr Atheroscler Rep 23, 22 (2021).

112. P. Pati et al., Time-restricted feeding reduces cardiovascular disease risk in obese mice. JCI Insight 10, (2025).

113. M. Milan et al., Time-restricted feeding improves aortic endothelial relaxation by enhancing mitochondrial function and attenuating oxidative stress in aged mice. Redox Biol 73, 103189 (2024).

114. R. A. Miller et al., Glycine supplementation extends lifespan of male and female mice. Aging Cell 18, e12953 (2019).

115. N. Ogiso et al., Biological characteristics of age-related changes in C57BL/6 mice sub-strains in the national center for geriatrics and gerontology aging farm. Exp Anim 74, 229–238 (2025).

116. W. J. Schwartz, P. Zimmerman, Circadian timekeeping in BALB/c and C57BL/6 inbred mouse strains. J Neurosci 10, 3685–3694 (1990).

117. M. Pfeffer, C. von Gall, H. Wicht, H. W. Korf, The Role of the Melatoninergic System in Circadian and Seasonal Rhythms-Insights From Different Mouse Strains. Front Physiol 13, 883637 (2022).

118. G. Banks et al., Genetic background influences age-related decline in visual and nonvisual retinal responses, circadian rhythms, and sleep. Neurobiol Aging 36, 380–393 (2015).

119. S. M. Siepka, J. S. Takahashi, Methods to record circadian rhythm wheel running activity in mice. Methods Enzymol 393, 230–239 (2005).

120. A. A. Bachmanov, D. R. Reed, G. K. Beauchamp, M. G. Tordoff, Food intake, water intake, and drinking spout side preference of 28 mouse strains. Behav Genet 32, 435–443 (2002).

121. D. R. Reed, A. A. Bachmanov, M. G. Tordoff, Forty mouse strain survey of body composition. Physiol Behav 91, 593–600 (2007).

122. J. W. Cooley, J. W. Tukey, An Algorithm for the Machine Calculation of Complex Fourier Series. Mathematics of Computation 19, 297–301 (1965).

123. R. B. Blackman, J. W. Tukey, The measurement of power spectra from the point of view of communications engineering — Part I. The Bell System Technical Journal 37, 185–282 (1958).

124. C. Wang, Q. Li, D. T. Redden, R. Weindruch, D. B. Allison, Statistical methods for testing effects on “maximum lifespan”. Mech Ageing Dev 125, 629–632 (2004).

